# Resistome diversity in bovine clinical mastitis microbiome, a signature concurrence

**DOI:** 10.1101/829283

**Authors:** M. Nazmul Hoque, Arif Istiaq, Rebecca A. Clement, Keylie M. Gibson, Otun Saha, Ovinu Kibria Islam, Ruhshan Ahmed Abir, Munawar Sultana, AMAM Zonaed Siddiki, Keith A. Crandall, M. Anwar Hossain

## Abstract

The bovine clinical mastitis (CM) milk is a large reservoir for diverse groups of resistomes, which play important roles in the pathogenesis of mastitis, but little is known about the concurrence of CM microbiome signature and its associated resistomes. Here we deciphered the total resistance (antibiotics and metals resistance, biofilm formation, quorum sensing) present in CM microbiome using whole metagenome sequencing (WMS) and *in vitro* cultural approaches. Significant correlation (*p*=0.001) was found between the resistome diversity and microbiome signature. We identified the strain-level microbiome diversity in four cattle breeds, with microbiome composition represented by the phyla *Proteobacteria*, *Bacteroidetes*, *Firmicutes*, *Actinobacteria* and *Fusobacteria* (contributing to >95.0% of total strains). However, the resistome diversity did not vary significantly (*p*=0.692) across the microbiomes of cattle breeds. The *in vitro* investigation showed that biofilm producing CM pathogens were resistant to most of the conventional antibiotics used for CM treatment, whereas these pathogens remained sensitive to five heavy metals (Cr, Co, Ni, Cu, Zn) at varying concentrations. We also found association of some genomic functional potentials such as bacterial flagellar movement and chemotaxis, regulation and cell signaling, phages-prophages, transposable elements, plasmids and oxidative stress in the pathophysiology of bovine CM. These findings of rapid and reliable identification of CM microbiomes and associated resistomes will help improve the optimization of therapeutic schemes involving antibiotics and metals usage in the prevention and control programs of bovine CM.

## Introduction

Mastitis is the foremost production and major economic burden confronted by the global dairy industry^1–3^. Bovine clinical mastitis (CM) is of special concern for milk producers in developing countries like Bangladesh, where dairying plays a pivotal role in the national economy. The CM milk from dairy animals is now considered to host a complex microbial community with great diversity^2–4^. The most frequently isolated pathogens are *Staphylococcus aureus, Escherichia coli, Klebsiella* spp., *Streptococcus* spp., *Mycoplasma* spp., *Enterobacter* spp., *Bacillus* spp., *Corynebacterium* species^5–8^. Therefore, accurate identification of pathogens causing CM enables appropriate choices for antimicrobial treatment and preventive mastitis management^8–10^. Over the past two decades, a wide range of phenotyping and genotyping methods have been implemented to study mastitis-causing bacteria^6–9^. Although culture-based techniques are in the forefront of detecting CM bacteria, these methods are time-consuming and have inherent drawback of not being applicable to non-cultivable bacteria^11^. Until recently, 16S rRNA partial gene sequencing remained as the most commonly used genomic survey tool to study bovine mastitis microbiomes^3,4,12^. However, this technique has limitations because of polymerase chain reaction (PCR) bias, lower taxonomic resolution at the species level, and limiting information on gene abundance and functional profiling^13^. Shotgun whole metagenome sequencing (WMS), on the other hand, produces a metagenome reflecting the breadth of microbial genomic content in a sample and successfully provides insights into the phylogenetic composition, species and/or strain and functional diversity for a variety of biomes^2,13,14^. This WMS typically produces high complexity datasets with millions of short reads allowing extensive characterization of microbiome in an ecological niche^13,14^ and profiling of their functional attributes like microbial energy metabolism, antimicrobial resistance and biofilm forming abilities; and gradually becoming a cost-effective metagenomic approach^13^. The cattle breeds or host genetics may have an influence on the milk microbiota composition and on susceptibility to disease and resistance to bacterial infection^12,15^. The milk from healthy Holstein Friesian cows displayed more significant changes bacterial biodiversity and composition than microbiota in Rendena cows milk^12,16^.

The secretion of antimicrobial compounds by microbes is an ancient and effective method to improve the survival of microbes competing for space and nutrients with other microorganisms^17^. However, the advent recent metagenomic studies have revealed diverse homologues of known resistance genes broadly distributed across environmental locales including bovine milk samples. This widespread dissemination of antimicrobial resistance elements is inconsistent with a hypothesis of contemporary emergence and instead suggests a richer natural history of resistance^18^. The vast diversity of bacterial species in CM milk coupled with short generation times and horizontal gene transfer permit the rapid accumulation of countless resistance variations at a relatively high evolutionary pace^19^. Resistance in CM bacteria typically goes unnoticed until a given species becomes of clinical interest, and the resistome found CM is also suspected to be a source of newly emerging resistance genes in the CM^2,8,17,20^. Antibiotics have been used for decades in livestock production for both therapeutic (e.g. treatment of specific diseases) and nontherapeutic (growth promotion) purposes^10^. However, there are data that support the fact that both nontherapeutic and therapeutic doses of antibiotics can contribute to the emergence of antimicrobial-resistant bacteria, thus exacerbating the problem of antibiotic resistance in animal and human pathogens^10^, and enhancing the selection for antibiotic resistance genes (ARGs) and the horizontal transfer of these genes^10,17^. Bacteria residing in the bovine gastrointestinal tract and udder may become resistant to these antibiotics and, once released into the milk, they may transfer ARGs to other CM bacteria of contagious and environmental origin^8,20^. Efficacy of antimicrobial therapy against bovine CM pathogens is low^8^, and the use of antibiotics, confined to selected severe CM cases necessitates the accurate identification and characterization of pathogens and antibiotic selection for its better prevention and control^1,8^. Furthermore, antimicrobial resistance (AMR) is a global health concern in both human and veterinary medicine^10^, and thus, monitoring the emergence of AMR bacterial strains is an essential component of bovine CM prevention and control strategies^8,21^. Therefore, finding an effective alternative strategy for the control of bovine mastitis is a challenge for dairy producers.

The antimicrobial properties of metals have been documented throughout the history of medicine and healthcare^22^. The metal salts such as chromium (Cr), cobalt (Co), nickel (Ni), copper (Cu) and zinc (Zn) are effective in controlling bacterial transmission and infection risks^22^. However, their uses are limited due to their toxicity and possible detrimental environmental effects in dairy industries particularly as therapeutic agents against bovine CM pathogens. Biofilm formation is an important virulence factor for mastitis causing bacteria and contributes to the resistance to different classes of antimicrobials^23^. Bacterial pathogens identified in this study showed broad spectrum of antimicrobial (antibiotics, toxic metals) resistance, and possessed biofilm forming and quorum sensing abilities, which might be the potential factors hindering CM cures, thereby leading to the persistence of the disease, and increased risk of transmission to non-infected dairy cows. Genetic information about resistance or *in vitro* assays of resistance is not enough to understand about resistomes when considered solely rather in combination^10,11^. Genetic potential doesn’t give the idea of resistance level as many other factors are involve such as expression, stimulation, stress etc^10,11,15^. Similarly, resistance assay doesn’t give the idea about genetic makeup responsible. Therefore, our present study describes the resistome diversity across microbial communities causing CM in four major cattle breeds (Local Zebu, LZ; Red Chattogram Cattle, RCC; Sahiwal, SW; Crossbred Holstein Friesian; XHF) of Bangladesh using both metagenomic deep sequencing (WMS) and *in vitro* cultural approaches. Furthermore, we also aimed to investigate the influences of metabolic genomic potentials of the microbiomes in the pathophysiology of bovine CM.

## Results

To decipher the resistome diversity in bovine CM microbiomes, we used a condition of combination of *in silico* (WMS, 16S rRNA gene sequencing) and *in vitro* (culture base) approaches. The present WMS investigation leads to the direct and comprehensive evaluation of resistance to antibiotics and toxic compounds (RATC), biofilm formation (BF) and quorum sensing (QS) genes in 25 CM samples. Furthermore, *in vitro* antimicrobial resistance profiling of six CM causing bacteria (*S. aureus*, *E. coli*, *Klebsiella*, *Enterobacter*, *Bacillus* and *Shigella*) isolated from 260 milk samples was carried out using 12 commonly used antibiotics (ampicillin, doxycycline, tetracycline, nitrofurantoin, ciprofloxacin, nalidixic acid, cefoxitin, imipenem, chloramphenicol, gentamycin, erythromycin, vancomycin), and five toxic metals (copper, zinc, chromium, nickel, cobalt). Moreover, we also demonstrated some functional metabolic potentials of CM microbiomes found to be associated with mammary gland pathogenesis.

### Sequence analysis

The WMS of 25 CM milk samples generated approximately 600 million reads, ranging from 8.86 to 39.75 million per sample. An average of 21.13 million reads per sample (maximum=36.89 million, minimum=4.71 million) passed the quality control step (Supplementary Data 1). We analyzed the sequencing reads simultaneously using two bioinformatics pipelines, PathoScope 2.0 (PS) and MG-RAST (MR).

### Microbiome diversity and composition in CM

We investigated the strain-level microbial community and relative abundances in 25 CM milk samples (previously published 14 samples^2^ and 11 new samples) through WMS. The reads generated from WMS mapped to 391 genera and 519 strains of bacteria through MR and PS analyses, respectively (Supplementary Data 1).

The rarefaction curves based on observed species richness reached a plateau after, on average, 23.87 million reads (Fig. 1a, Supplementary Data 1)-suggesting that the depth of coverage for most samples was sufficient to capture the entire microbial diversity within each sample. Although, we did not find any significant differences in the alpha (observed species, Chao1, ACE, Shannon, Simpson and Fisher diversity estimates) and beta (based on Bray-Curtis dissimilarity matrix) diversities among the microbial communities across the 25 CM samples (Fig. 1b,c). However, significant diversity (alpha and beta) differences were observed among the CM microbiome communities across the four cattle breeds (LZ, RCC, SW, XHF) regardless of the method (i.e., either PS or MR) used to tabulate microbial abundances (PS; *p*=0.005, MR; *p*=0.001, Kruskal–Wallis test). In addition, this breed specific diversity difference remained evident in the microbial ecosystem of XHF cows associated CM milk samples (Fig. 1d,e). The PCoA analysis also showed significant microbial disparity (*p*=0.001) among the microbiome of four dairy breeds (Fig. 1e).

**Fig. 1.**
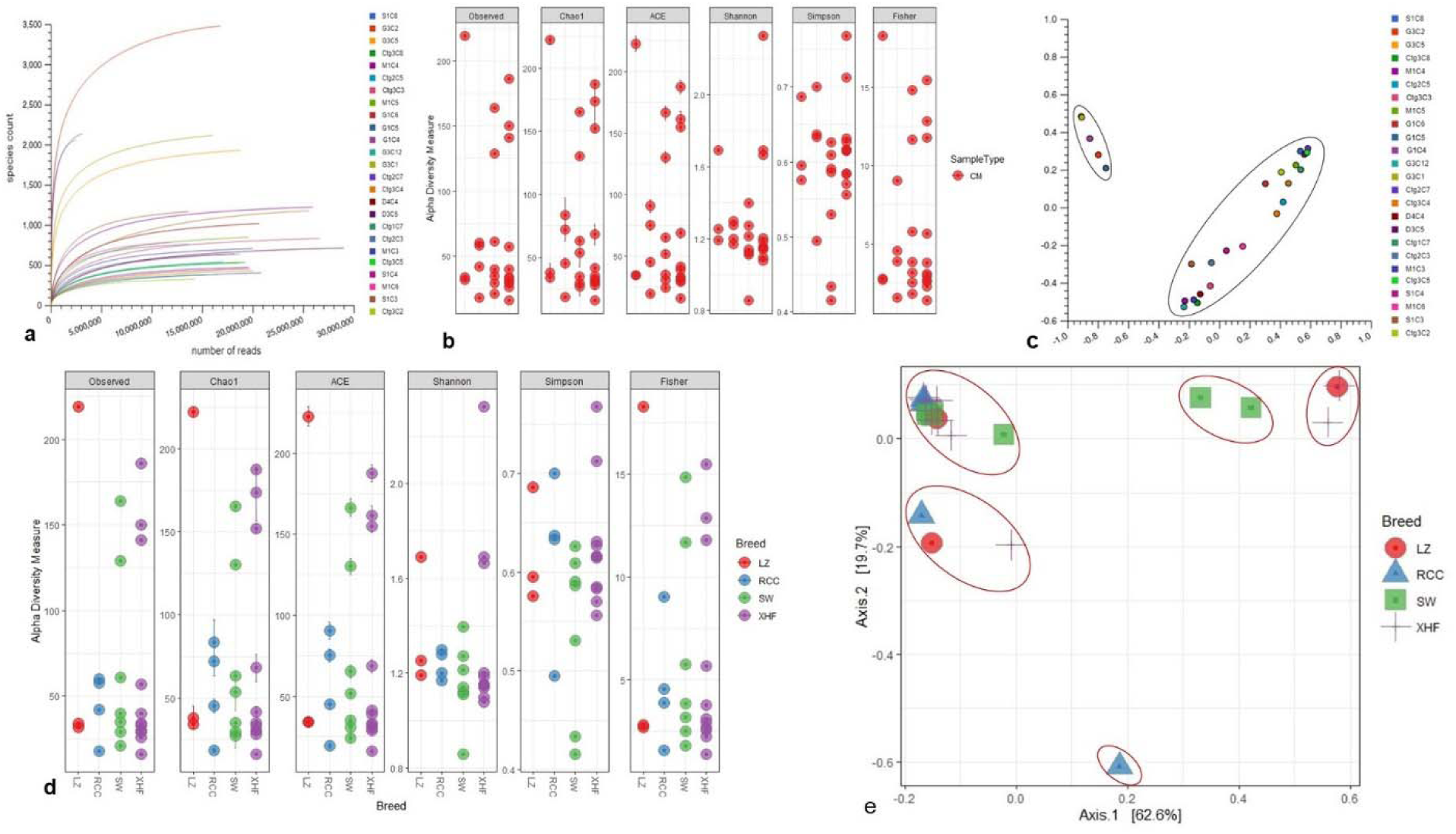
Bovine clinical mastitis (CM) milk microbiome diversity. **a)** Rarefaction curves showing the influence of sequencing depth (number of reads per sample, X axis) on species richness (Y axis) in CM milk samples. The rarefaction curves representing the number of species per sample indicated that the sequencing depth was sufficient enough to fully capture the microbial diversity as existed. **b)** Alpha diversity measured using the observed species, Chao 1, ACE and Shannon diversity indices through PathoScope (PS) analysis. The observed species richness (P_Observed_ = 0.511), Chao1 (P_Chao1_ = 0.081), ACE (P_ACE_ = 0.121), Shannon (P_Shannon_ = 0.401), Simpson (P_Simpson_ = 0.011) and Fisher (P_Fisher_ = 0.014) diversity analyses revealed that microbiome diversity did not vary among the CM samples. **c)** Beta diversity (Principal coordinate analysis; PCoA) measured on the Bray-Curtis distance method using MG-RAST tool for CM causing microbial communities (genus-level) shows that most of the CM samples clustered together (black circle) indicating no significant diversity differences. **d)** Alpha diversity measured using species richness (P_Observed_ = 0.011), Chao1 (P_Chao1_ = 0.001), ACE (P_ACE_ = 0.021), Shannon (P_Shannon_ = 0.001), Simpson (P_Simpson_ = 0.009) and Fisher (P_Fisher_ = 0.023) diversity matrices on PS data showed significant diversity differences (Kruskal–Wallis test, *p*=0.002) within the microbial communities of four breeds (Local Zebu cows, LZ; Red Chattogram cows, RCC; Sahiwal, SW; Holstein Friesian cross, XHF) of cows. **e)** PCoA plot based on weighted-UniFrac distance method at strain-level microbiome signature of four breeds of cows reveals that the CM samples appear more distantly (red circles) indicating significant group differences (*p*=0.001). This differences in the microbiome signature associated with CM in four breeds could be explained by a large percentage of variation in the first (62.6%) and second (19.7%) axes.

The predominant bacterial phyla were *Proteobacteria*, *Bacteroidetes*, *Firmicutes*, *Actinobacteria* and *Fusobacteria* (contributing to >95.0% of the total sequences, Kruskal–Wallis test, *p*=0.001) in the MR analysis. The strain-level signature of the microbiome demonstrated that most of the species identified in each CM sample represented by multiple strains (Supplementary Data 1), and of the detected bacterial strains, the top 200 strains (according to their relative abundance) are depicted in Fig. 2. The CM associated microbiome was dominated by 29 different strains of *Pseudomonas* species, while *Acinetobacter*, *Streptococcus*, *Lactobacillus*, *Corynebacterium*, *Staphylococcus* and *Enterococcus* species represented by 27, 27, 18, 17, 15 and 10 different strains, respectively (Fig. 2, Supplementary Data 1). Thus, among the identified bacterial strains, *A. johnsonii* XBB1 had the highest relative abundance (38.9%) and followed by *Micromonospora* sp. HK10 (17.6%). Other bacterial strains found abundantly were *Campylobacter mucosalis* (8.7%), *P. putida* KT2440 (7.7%), *Anaerobutyricum hallii* DSM 3353 (6.3%), *P. fragi* (3.2%), *Catenibacterium mitsuokai* DSM 15897 (3.0%), *E. coli* O104:H4 str. 2011C-3493 (2.0%), *A. veronii* (1.2%), *Pantoea dispersa* EGD-AAK13 (1.1%), *P. fluorescens* Pf0-1 (0.8%), *K. oxytoca* (0.7%) and *P. entomophila* L48 (0.5%). The remaining strains had a relatively lower abundance (<0.5%) (Supplementary Data 1). According to the cattle breeds, the XHF cows had the highest number of microbial strains (n=403) followed by LZ cows (n=230), SW cows (n=134) and RCC (n=125) (Fig. 3a-c, Supplementary Data 1). The breed specific association revealed that 45.7, 22.6 and 19.1% of the detected bacterial strains in CM milk samples of LZ, SW and RCC cows, respectively, were also found in the CM microbiome of XHF cows (Fig. 3d, Supplementary Data 1).

**Fig. 2.**
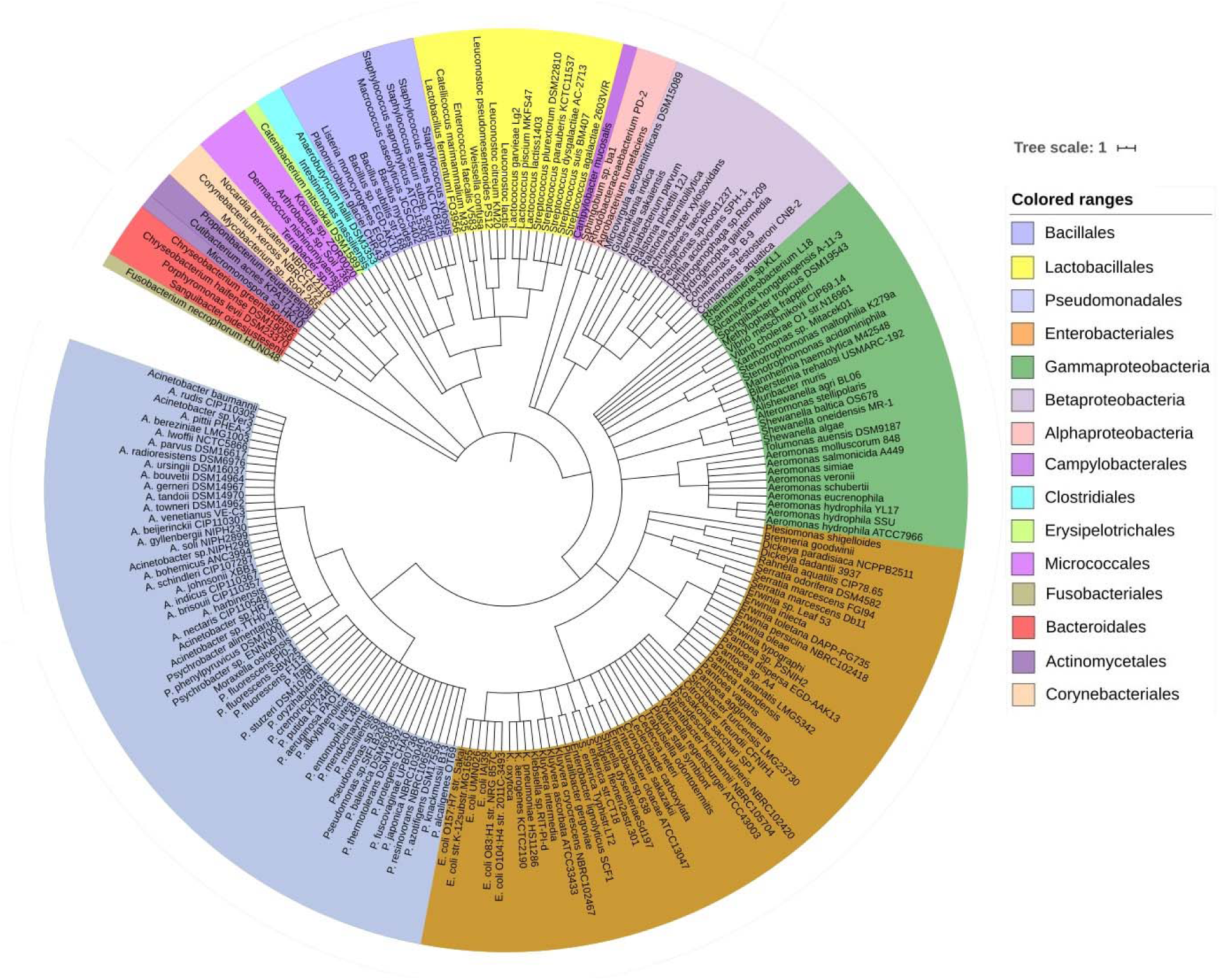
The strain-level taxonomic profile microbiota associated with bovine clinical mastitis (CM). Taxonomic dendrogram showing the top bacterial microbiome of bovine CM milk. Color ranges identify different strains within the tree. Taxonomic dendrogram was generated with the top 200 abundant unique strains of bacteria in CM milk metagenome based on the maximum likelihood method in Clustal W and displayed with iTOL (interactive Tree Of Life). Each node represents a single strain shared among more than 50 % of the samples at a relative abundance of >0.0006% of the total bacterial community. The inner circle represents the root of the microbiome defined as bacteria present in 25 CM milk samples. The outer circle shows the strains and/or species colored by different order of bacteria present in >80% of samples. The strains in the phylogenetic tree are also available in Supplementary Data 1.

**Fig. 3.**
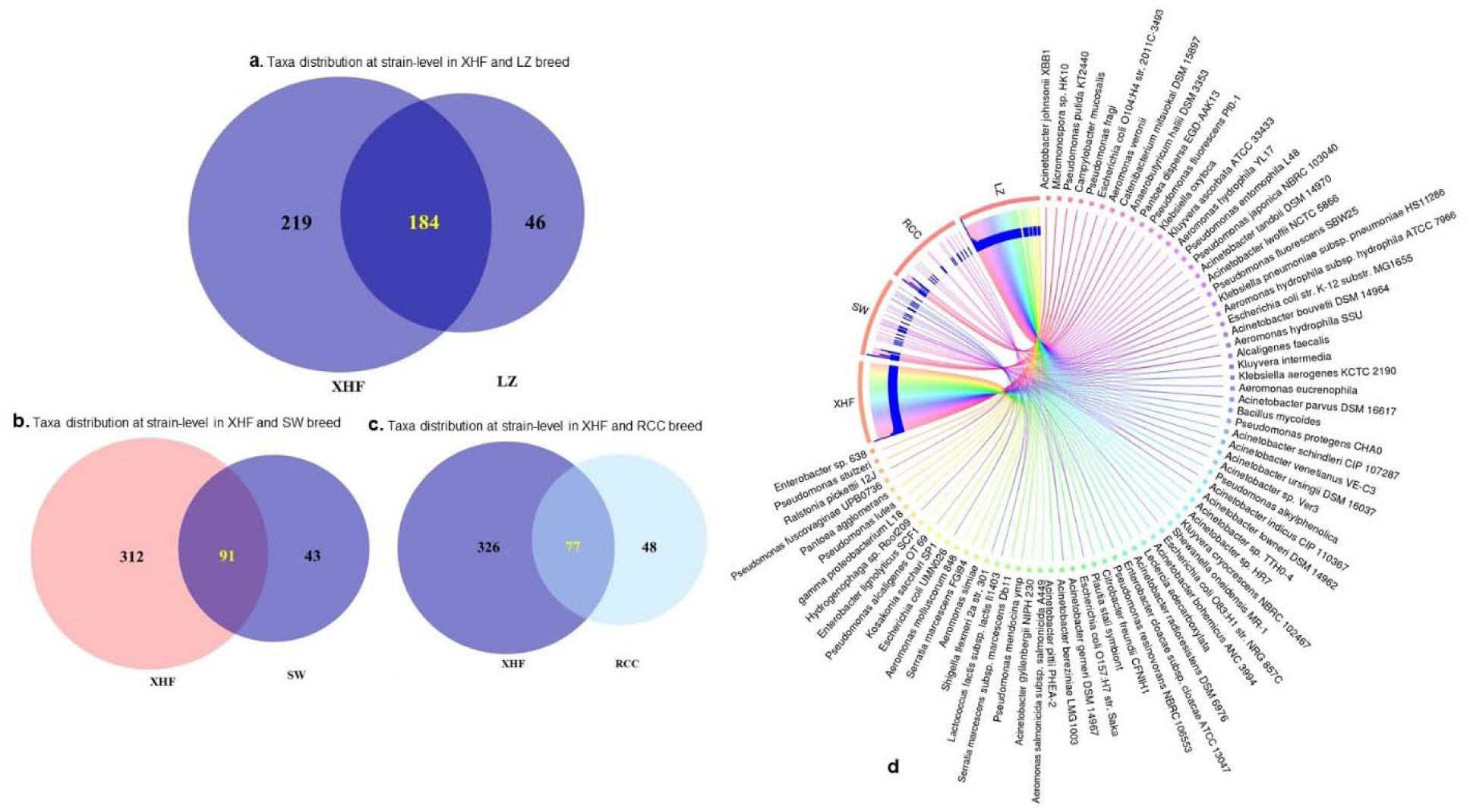
Strain-level bovine CM microbiome diversity in four different breeds (Local Zebu, LZ; Red Chattogram Cattle, RCC; Sahiwal, SW; Crossbred Holstein Friesian, XHF) of cows through PathoScope (PS) analysis. **a)** Venn diagrams representing the core unique and shared microbiomes of bovine clinical mastitis (CM) in XHF and LZ breeds while **b)** and **c)** Venn diagrams showing the unique and shared bacterial strains in XHF and SW and XHF and RCC breeds, respectively. Microbiome sharing between the conditions are indicated by yellow color. **d)** The circular plot illustrates the relative abundance of the top 75 CM causing bacterial strains in CM milk samples obtained from XHF, LZ, SW and RCC dairy breeds. Taxa in the respective breed of cows are represented by different colored ribbons, and the inner blue bars indicate their respective relative abundances. The XHF cows had the highest number of microbial strains followed by LZ, SW and RCC. This breed specific association revealed that 45.66, 22.58 and 19.11% of the detected bacterial strains in CM milk collected from LZ, SW and RCC cows, respectively, were also seen in the CM milk microbiome of XHF cows. The relative abundance bacterial strains in four breeds is also available in Supplementary Data 1.

Simultaneously through *in vitro* cultural analysis, a total of 452 isolates that belonged to six bacterial (*S. aureus*, *E. coli*, *Klebsiella*, *Enterobacter*, *Bacillus* and *Shigella*) species were identified in 260 CM samples (including 25 WMS CM samples) collected from central (CR=160) and southeastern (SER=100) regions of Bangladesh (Supplementary Fig. 1). The overall prevalence of *S. aureus*, *E. coli*, *Klebsiella*, *Enterobacter*, *Bacillus* and *Shigella* species were 23.5, 18.5, 19.2, 12.3, 9.2 and 17.3% CM samples, respectively (Supplementary Table 1). We found significant differences in the prevalence of these species (*p*=0.01) when analyzing the distribution of these pathogens according to the origin of the samples (SER and CR) (Supplementary Fig. 2). The culture-based findings of the current study demonstrated *S. aureus* as the chief etiology of bovine CM in Bangladesh, while *Shigella* species remained as the least frequently detected CM pathogen – which corroborates with the results of WMS-based taxonomic identification (Supplementary Fig. 3).

### Resistomes diversity and composition of CM microbiome

For analyses of resistome diversity and abundance in CM microbiomes, the SEED module of the MR pipeline provided a comprehensive picture. Using SEED, 147,040 reads aligned to 30 resistance to antibiotics and toxic compounds (RATC) and 10 biofilm formation and quorum sensing (BF-QS) functional groups across the CM samples with different abundances (Supplementary Data 2). The RATC genes classified into two unique groups, 19 antibiotic resistance and 11 toxic metal resistance groups (Fig. 4, Supplementary Data 2). This WMS analysis showed significant correlation (Pearson correlation, *p*=0.001; Nonparametric Spearman’s Correlation, *p*=0.003) between the number of reads aligned to bacterial genomes and number of reads mapped to RATC genes (Supplementary Data 2). Among the RATC functional groups, multidrug resistance to efflux pumps (MREP, 28.6%), *Cme*ABC operon (8.9%), resistance to fluoroquinolones (RFL, 6.2%), *mdt*ABCD cluster (5.5%), methicillin resistance in *Staphylococci* (MRS, 3.8%), *Bla*R1 regulatory family (*Bla*R1, 3.4%), *Mex*E-*Mex*F-*Opr*N (2.4%) and beta-lactamase resistance (ΒLAC, 2.2%) were the dominating antibiotic resistance genes (ARGs) found in CM milk microbiomes (Fig. 4a, Supplementary Data 2). In addition to ARGs, the WMS analysis also detected a number of metal and toxic compound resistance (MTR) genes in CM microbiomes. Among them, cobalt-zinc-cadmium resistance (CZCR, 19.3%), copper homeostasis (CH, 9.6%), arsenic resistance (AR, 2.9%), copper homeostasis: copper tolerance (CHCT, 2.3%) and resistance to chromium compounds (RCHC, 1.4%) were the predominating resistant genes (Fig. 4a, Supplementary Data 2). Although the relative abundance of these RATC genes varied among the microbiomes of the four breeds (LZ, RCC, SW and XHF), but their resistome diversity did not vary significantly (*p*=0.692) by taxonomic diversity of respective breeds (Fig. 4b, Supplementary Data 2).

**Fig. 4.**
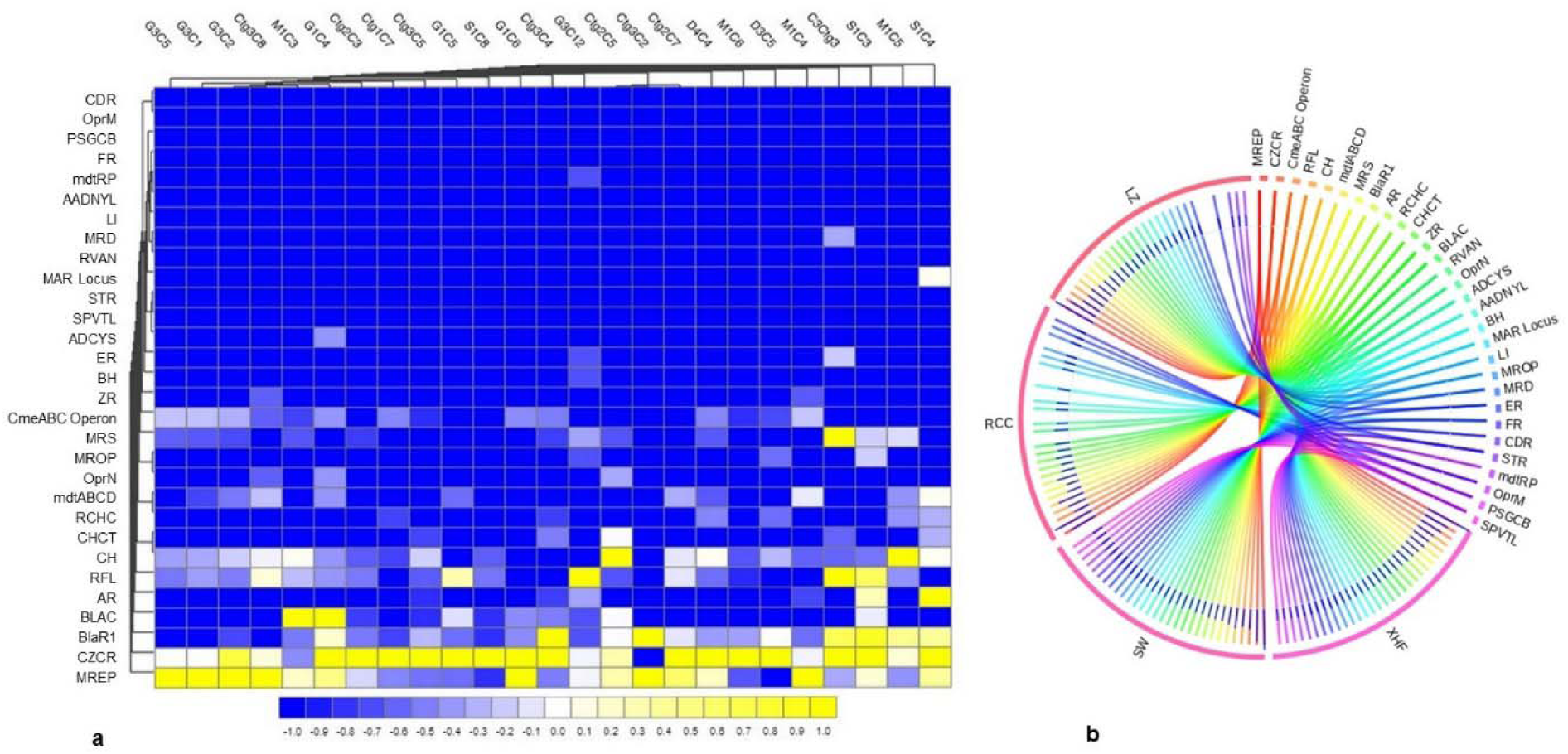
Projection of the resistance to antibiotic and toxic compounds (RATC) genes in bovine clinical mastitis (CM) pathogens. **a)** Heatmap showing the hierarchical clustering of 30 different RATC genes detected in CM associated microbiomes of 25 CM milk samples as measured at level-3 of SEED subsystems in MG-RAST pipeline. The relative abundance of these genes significantly correlated (Pearson correlation, *p*=0.002) with the relative abundance of the bacterial taxa found in these samples. The color bar at the bottom represents the relative abundance of putative genes and expressed as a value between −1 (low abundance) and 1 (high abundance). The yellow color indicates the more abundant patterns, while blue cells for less abundant RATC gene in that particular sample. **b)** The circular plot illustrates the diversity and relative abundance of the RATC genes detected among the microbiomes of the four different breeds (Local Zebu, LZ; Red Chattogram Cattle, RCC; Sahiwal, SW; Crossbred Holstein Friesian, XHF) of cows through SEED subsystems analysis. We found no significant correlation between the resistome and microbiome diversity in different breeds (*p*=0.692). The association of the RATC genes according to breeds is shown by different colored ribbons and the relative abundances these genes are represented by inner blue colored bars. Part of the RATC functional groups are shared among microbes of the four breeds (XHF, LZ, SW and RCC), and some are effectively undetected in the microbiomes of the other breeds. Abbreviations: MREP, multidrug resistance efflux pumps; CZCR, cobalt-zinc-cadmium resistance; BlaR, BlaR1 family regulatory sensor-transducer disambiguation; BLAC, beta-lactamase resistance; AR, arsenic resistance; RFL, resistance to fluoroquinolones; CH, copper homeostasis; CHCT, copper homeostasis: copper tolerance; RCHC, resistance to chromium compounds; mdtABCD, the mdtABCD multidrug resistance cluster; OprN, mexe-mexf-oprn multidrug efflux system; MROP, mercury resistance to operon; MRS, methicillin resistance in *Staphylococci*; CmeABC Operon, multidrug efflux pump in *Campylobacter jejuni*; ZR, zinc resistance; BH, bile hydrolysis; ER, erythromycin resistance; ADCYS, adaptation to d-cysteine; SPVTL, *Streptococcus pneumoniae* vancomycin tolerance locus; STR, Streptothricin resistance; MAR Locus, multiple antibiotic resistance to locus; RVAN, resistance to vancomycin; MRD, mercuric reductase; LI, lysozyme inhibitors; AADNYL, aminoglycoside adenylyltransferases; mdtRP, multidrug resistance operon mdtRP of *Bacillus*; FR, Fosfomycin resistance; PSGCB, polymyxin synthetase gene cluster in *Bacillus*; OprM, mexA-mexB-oprm multidrug efflux system; CDR, cadmium resistance. Additional information is also available in Supplementary Data 2.

The diversity and composition of RATC functional groups also varied significantly (*p*=0.027) in *in vitro* selected six CM pathogens isolated and identified from different sources of CM samples (breed and study areas) under almost same farming management system (Fig. 5a, Supplementary Data 2). Among the RATC groups, the predominant ARGs found as follows MRS (*S. aureus*, 37.0%), RFL (*S. aureus*, 14.8%; *Shigella*, 7.8%), MREP (*E. coli*, 28.5%; *Klebsiella*, 28.4%), *Bla*R1 (*E. coli*, 6.0%; *Shigella*, 8.5%), *mdt*ABCD cluster (*E. coli*, 17.5%; *Klebsiella*,18.9%; *Enterobacter*, 21.4%; *Shigella*, 11.7%), multiple antibiotic resistance (MAR) Locus (*E. coli*, 2.4%; *Enterobacter*, 2.6%), *Cme*ABC operon (*E. coli*, 9.1%; *Enterobacter*, 11.0%; *Shigella*, 25.6%), and adaptation to d-cysteine, ADCYS (*Bacillus*, 5.5%) (Fig. 5b). Conversely, genes encoding CH in *S. aureus* (11.1%), *E. coli* (4.8%), *Enterobacter* (4.4%), and *Shigella* (6.0%), CHCT in *Klebsiella* (11.2%) and *Shigella* (3.7%), mercuric reductase (MRD) in *S. aureus* (11.1%), mercury resistance to operon (MROP) in *Enterobacter* (2.4%), AR in *S. aureus* (3.7%), *E. coli* (4.4%), *Klebsiella* (10.1%), *Enterobacter* (7.5%) and *Shigella* (7.8%), ZR in *E. coli* (5.6%), cadmium resistance (CDR) in *S. aureus* (3.7%), CZCR in *S. aureus* (3.7%), *E. coli* (10.4%), *Klebsiella* (11.6%), *Enterobacter* (20.3%) and *Shigella* (21.0%), and RCHC in *Bacillus* (85.0%) were the most abundant toxic compounds or metals resistant (MTR) RATC functional groups among the six selected pathogens (Fig. 5c). Assessment of the BF-QS ability of the CM microbiomes revealed that autoinducer 2 (AI-2) transport and processing (*lsr*ACDBFGE operon, 33.7%), biofilm adhesion biosynthesis (BAB, 24.2%), protein *Yjg*K cluster linked to biofilm formation (*Yjg*K cluster, 15.5%), quorum sensing: autoinducer-2 synthesis (QSAU2, 9.4%) were the most abundant genes among CM associated pathogens (Supplementary Data 2). However, by comparing the association of these BF-QS genes among the selected six bacterial pathogens, we found significant variation (*p*=0.017) in their diversity, composition and relative abundances (Fig. 5d, Supplementary Data 2).

**Fig. 5.**
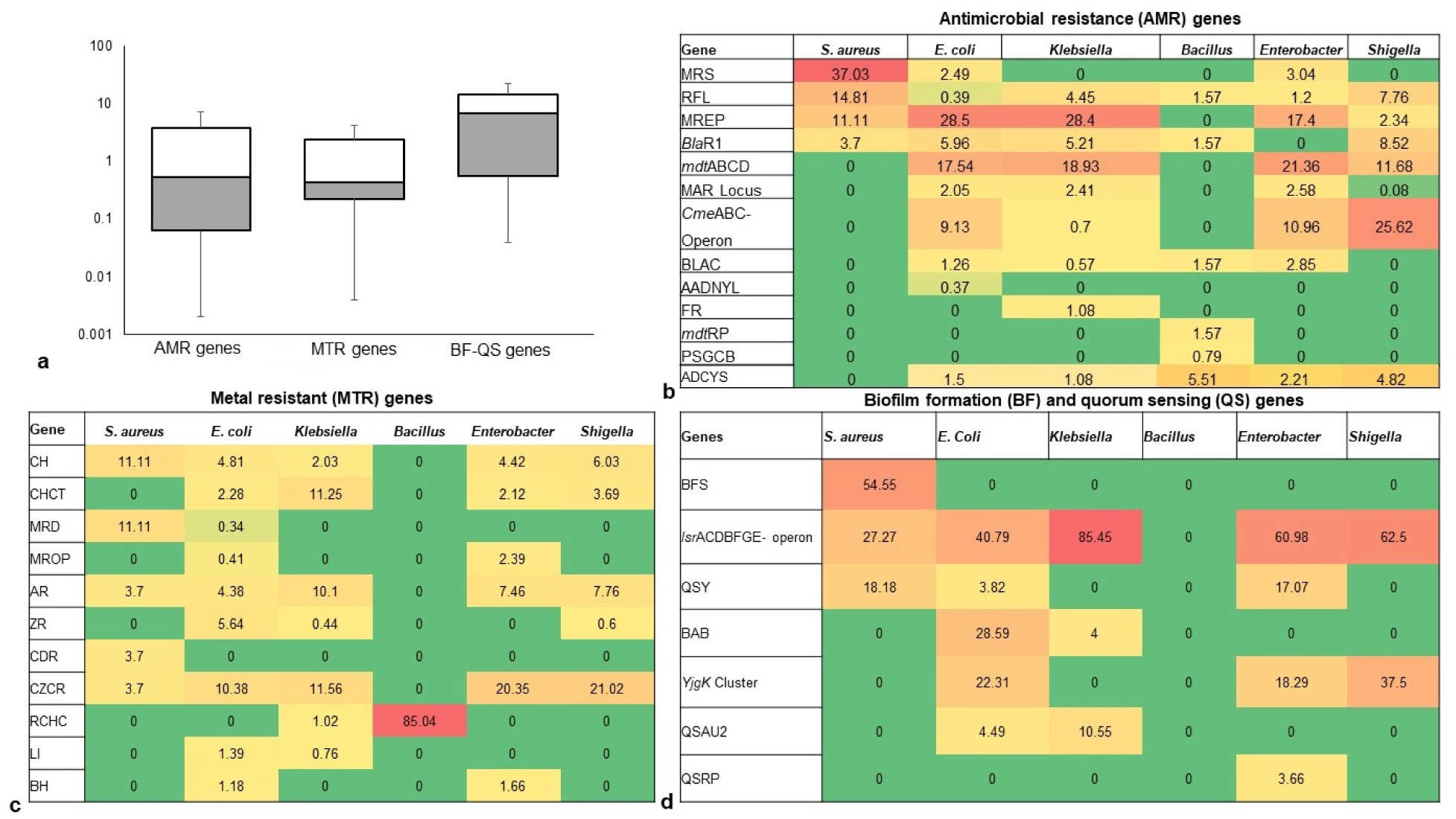
Heatmap comparison of antibiotics, metals, biofilm formation and quorum sensing genes found in the metagenome sequences (WMS) of six CM causing bacteria through SEED subsystems analysis in MG-RAST pipeline. **a)** Diversity and relative abundance of the antimicrobial resistance (AMR), metal resistance (MTR), and biofilm formation (BF) and quorum sensing (QS) genes varied significantly (Kruskal–Wallis test, *p*=0.029) among the study bacteria. **b)** Relative abundance of AMR genes, **c)** Relative abundance of MTR genes **d)** Relative abundance of BF-QS genes. Values are colored in shades of green to yellow to red, indicating low (absent), medium and high abundance, respectively. Abbreviations: MRS, methicillin resistance in *Staphylococci*; RFL, resistance to fluoroquinolones; MREP, multidrug resistance to efflux pumps; BlaR, BlaR1 family regulatory sensor-transducer disambiguation; mdtABCD, the mdtABCD multidrug resistance cluster; MAR Locus, multiple antibiotic resistance; CmeABC Operon, Multidrug efflux pump in *Campylobacter jejuni*; BLAC, beta-lactamase resistance; AADNYL, aminoglycoside adenylyltransferases (Gentamycin resistance); FR, Fosfomycin resistance; mdtRP, multidrug resistance operon mdtRP of *Bacillus*; PSGCB, polymyxin synthetase gene cluster in *Bacillus*; BFS, biofilm formation in *Staphylococcus*, lsrACDBFGE operon, autoinducer 2 (AI-2) transport and processing; QSY, quorum sensing in *Yersinia*; BAB, biofilm adhesion biosynthesis; *Yjg*K cluster, protein *Yjg*K cluster linked to biofilm formation; QSAU2, quorum sensing: autoinducer-2 synthesis; QSRP, quorum sensing regulation in *Pseudomonas*; CH, copper homeostasis; CHCT, copper homeostasis: copper tolerance; MRD, mercuric reductase; MROP, mercury resistance to operon; AR, arsenic resistance; ZR, zinc resistance; CDR, cadmium resistance; CZCR, cobalt-zinc-cadmium resistance; ADCYS, adaptation to d-cysteine; RCHC, resistance to chromium compounds; LI, lysozyme inhibitors; BH, bile hydrolysis. More details about these genes can be found in the text and Supplementary Data 2.

The *in vitro* antibiogram profiling of 221 individual isolates of the six bacteria revealed that *S. aureus* isolates had highest resistance to doxycycline, ampicillin, tetracycline and erythromycin (73.0 to 88.0%) and moderate resistance to chloramphenicol, ciprofloxacin and nitrofurantoin (50.0 to 58.0%) (Fig. 6, Table 1). The isolates of another Gram-positive bacterium (*Bacillus*) demonstrated highest resistance against doxycycline, ampicillin, nalidixic acid and erythromycin (60.0 to 84.0%). However, *E. coli* isolates exhibited highest resistance against tetracycline, doxycycline, nalidixic acid and ampicillin (77.0 to 93.0%) and moderate resistance to chloramphenicol, nitrofurantoin, gentamicin and ciprofloxacin (40.0 to 63.0%). The isolates of *Klebsiella*, *Enterobacter* and *Shigella* species displayed highest resistance to doxycycline, nalidixic acid, tetracycline and ampicillin (70.0 to 100.0%) and moderate resistance to ciprofloxacin, gentamicin, nitrofurantoin and chloramphenicol (30.0 to 70.0%). In this study, imipenem and cefoxitin remained as the most sensitive antibiotics against four Gram-negative bacterial (*E. coli*, *Klebsiella*, *Enterobacter* and *Shigella*) species, while the two Gram-positive (*S. aureus* and *Bacillus*) species were mostly sensitive to imipenem, cefoxitin and vancomycin (Fig. 6, Table 1). Taken together, the antibiogram profile revealed that all of the selected CM pathogens are becoming multidrug resistant (MDR, resistant to ≥5 antibiotics) and the highest resistance was found to tetracyclines (tetracycline and doxycycline) followed by quinolones (nalidixic acid) and penicillin (ampicillin) groups of antibiotics (Fig. 6, Table 1).

**Fig. 6.**
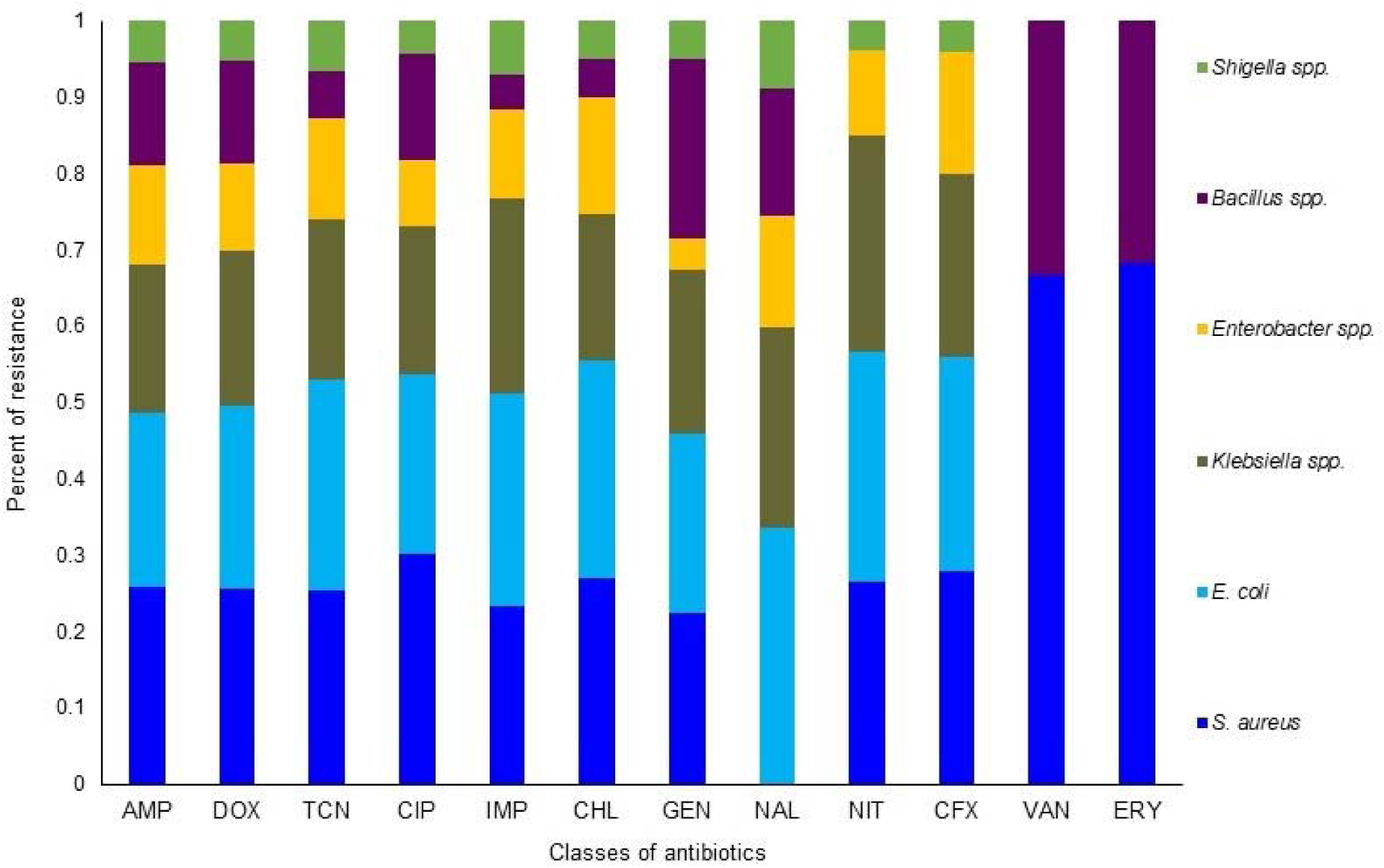
Antibiotic resistance pattern of bovine clinical mastitis pathogens by disk diffusion method. The antimicrobial resistance (AMR) patterns of the six bacteria obtained from 221 CM isolates (*S. aureus*, 56; *E. coli*, 54; *Klebsiella* spp., 42; *Enterobacter* spp., 26; *Bacillus* spp., 31; *Shigella* spp., 12) for twelve commonly used antibiotics from nine different groups/classes. Abbreviations: AMP, Ampicillin; DOX, Doxycycline; TCN, Tetracycline; CIP, Ciprofloxacin; IMP, Imipenem; CHL, Chloramphenicol; GEN, Gentamycin; NAL, Nalidixic acid; NIT, Nitrofurantoin; CFX, Cefoxitin; VAN, Vancomycin; ERY, Erythromycin. More details about AMR profiles can be found in the text and in Table 1.

**Table 1:**
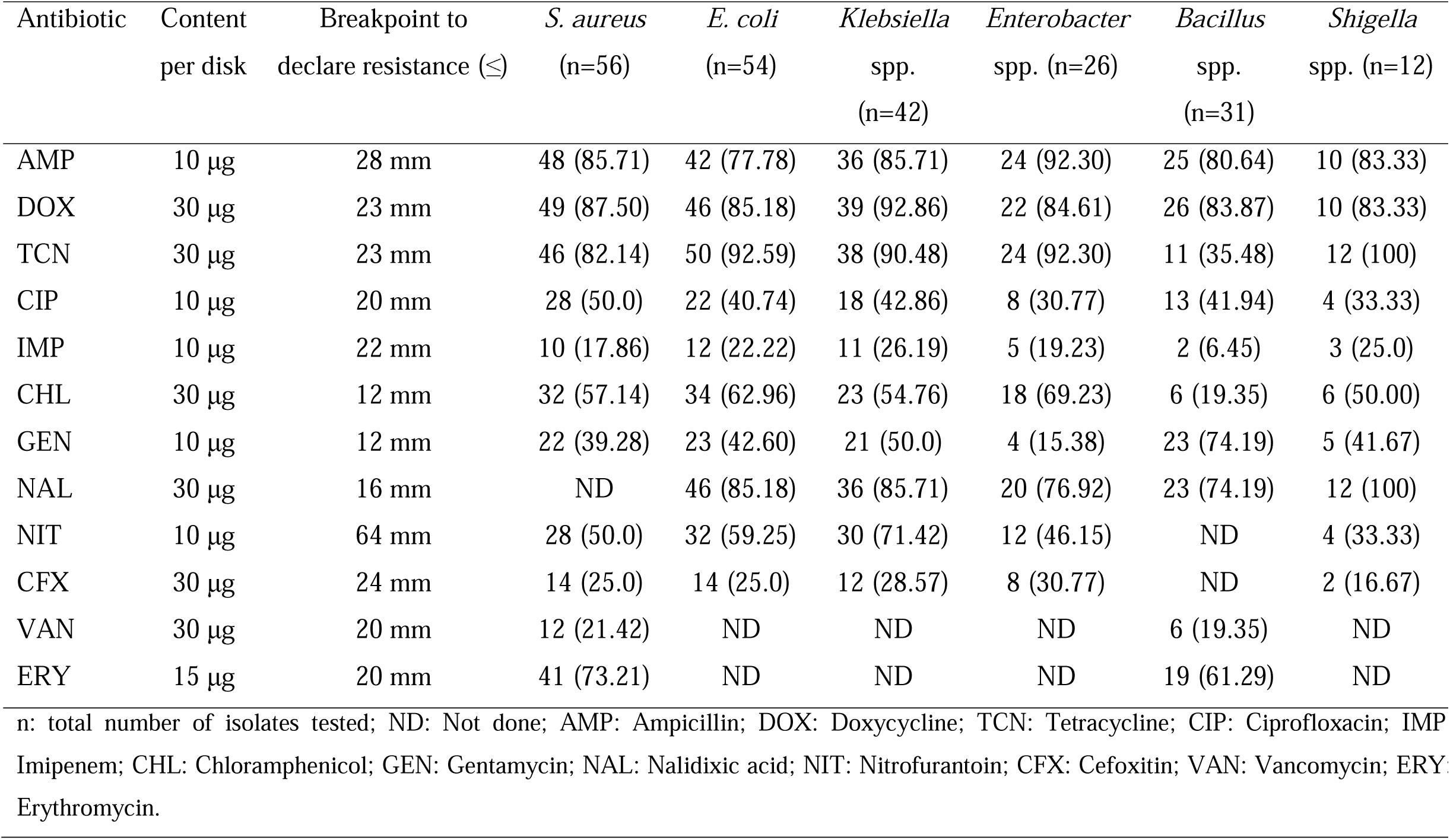
Antibiotic resistance pattern of bacteria [n (%) of isolates] associated with bovine clinical mastitis (CM).

The use of heavy metals in soluble forms as an alternative to prevent bovine CM appears as a novel promising idea supported by several earlier studies^1,22^. Zones of inhibition (ZOI) assays using the individual metal solution (Cu, Zn, Cr, Co and Ni) demonstrated an increase in antimicrobial activity which correlated with increased metal ion solution concentration (*p*<0.001) (Fig. 7). Thus, ZOI assays of metals demonstrated *S. aureus* (ZOI: 25.4 mm) as the most sensitive CM pathogens followed by *Bacillus* (ZOI: 23.4 mm), *E coli* (ZOI: 20.6 mm), *Enterobacter* (ZOI:18.9 mm), *Klebsiella* (ZOI:17.8 mm) and *Shigella* (ZOI:15.4 mm) species (Fig. 7a). The minimal inhibitory concentration (MIC) of the metal ions demonstrated a varying degree of response against all the tested CM pathogens, and these bacteria tolerated a wide range of metal concentration (3.4 to 38.1 μg/mL) (Supplementary Data 2). We compared the highest MIC values of each metal, and found that highest MIC values decrease in the following order: Zn (38.1 μg/mL, *S. aureus*), Cu (33.2 μg/mL, *Enterobacter* species), and Co μg/mL, *Bacillus* spp.) (Fig. 7b, Supplementary Data 2). For the MIC of specific bacteria, the most effective metals were found to be Cr against *Shigella* (3.4 μg/mL) and *Klebsiella* (5.8 μg/mL) species, Ni against *Shigella* (3.5 μg/mL) species, Co against *Shigella* (5 μg/mL) species, and Cu and Zn against *Shigella* (7.5 μg/mL, g/mL) and Cu (33.2 μg/mL) were the least toxic metals against *S. aureus* (Fig. 7b, Supplementary Data 2). A similar pattern was demonstrated for the minimal bactericidal concentration (MBC) with the greatest bactericidal activity for Cr against *S. aureus* (11.3 μg/mL) followed by Co against *E. coli* (14.3 μg/mL), Ni against *S. aureus* (23.1 μg/mL), Zn against *E. coli* (24.2 μg/mL), and Cu against *Shigella* (25.1 μg/mL) species. However, Cu produced equable antimicrobial efficacy as Zn, Cr, Co and Ni against *Enterobacter* species (≤25.5 μg/mL) (Supplementary Table 2).

**Fig. 7.**
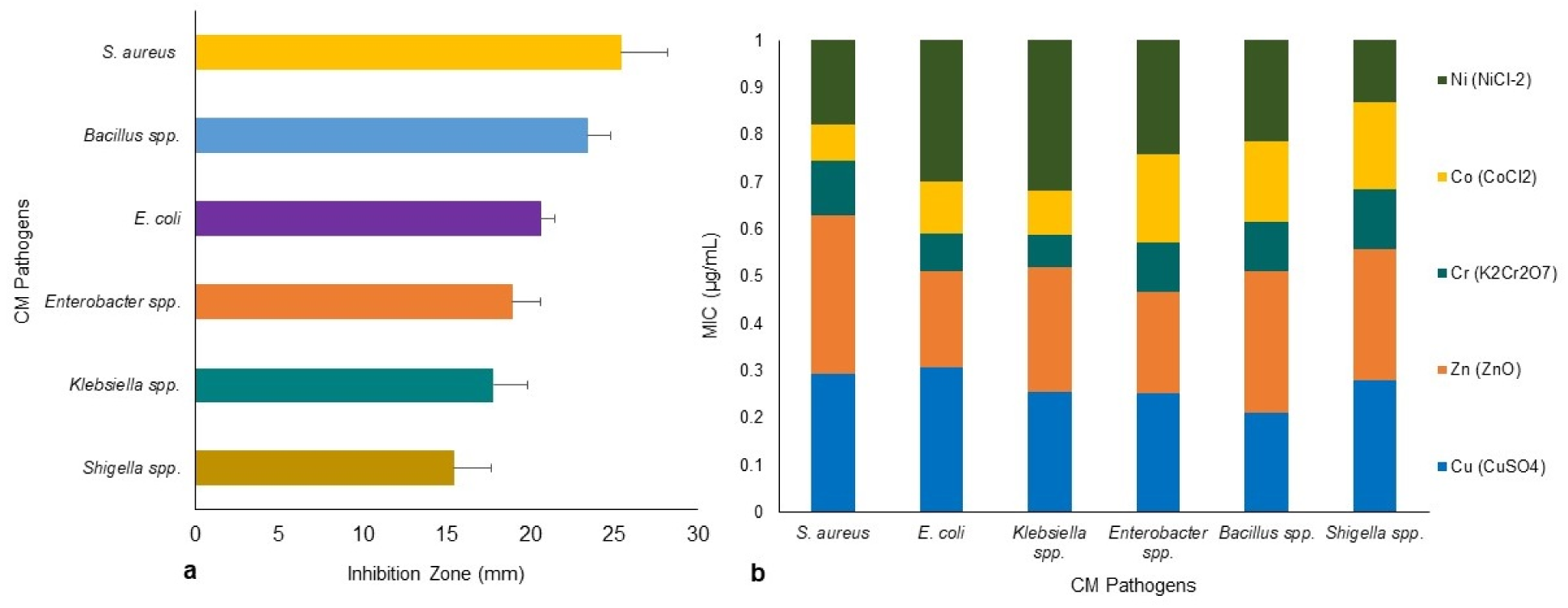
Antibacterial activity of heavy metals: Cu (CuSO4), Zn (ZnO), Cr (K2Cr2O7), Co (CoCl2) and Ni (NiCl2) against bovine CM pathogens. **a)** Zone of inhibition (ZOI, mm) for six CM causing bacteria, each bar representing the mean values (values given horizontal axis of the bars, mm) and standard deviation error bar (SD error bar) for each bacterium. **b)** Minimal inhibitory concentration (MIC) (expressed as μg/mL) of the tested metals against representative genera/species as determined by agar well diffusion and tube dilution methods.

To assess BF ability of CM pathogens in *in vitro* condition, we randomly selected 80 isolates (*S. aureus*, 15; *E. coli*, 15; *Klebsiella*, 15; *Bacillus*, 15; *Enterobacter*, 10 and *Shigella*, 10) for BF assay. In this study, 76.2% (61/80) bacterial species were biofilm producers with significance differences (*p*=0.028), and their categories of BF were strong biofilm forming (SBF, 28.7%), moderate biofilm forming (MBF, 25.2%), weak biofilm forming (WBF, 22.2%) and non-biofilm forming (NBF, 23.7%) (Fig. 8). While investigated individually, *E. coli* (66.7%) remained as the highest biofilm producing CM pathogen followed by *Enterobacter* (60.0%), *Klebsiella* (46.7%), *S. aureus* (40.0%), *Shigella* (30.0%) and *Bacillus* (26.7%) species. Our current findings revealed that Gram-negative CM pathogens (*Enterobacter*, 60.0%; *E. coli*, 40.0%; *Shigella*, 33.3%; *Klebsiella*, 28.6%) had higher biofilm producing ability than Gram-positive bacteria (*S. aureus*, 16.7%) (Fig. 8a,b). On the contrary, the majority of the *Bacillus* (73.3%), *Shigella* (70.0%) and *S. aureus* (60.0%) isolates remained as non-biofilm formers (NBF) (Fig. 8b). Therefore, our current findings of *in vitro* resistance analysis (antibiotics and metals resistance and biofilm assays) corroborate the resistome found in metagenome sequencing.

**Fig. 8.**
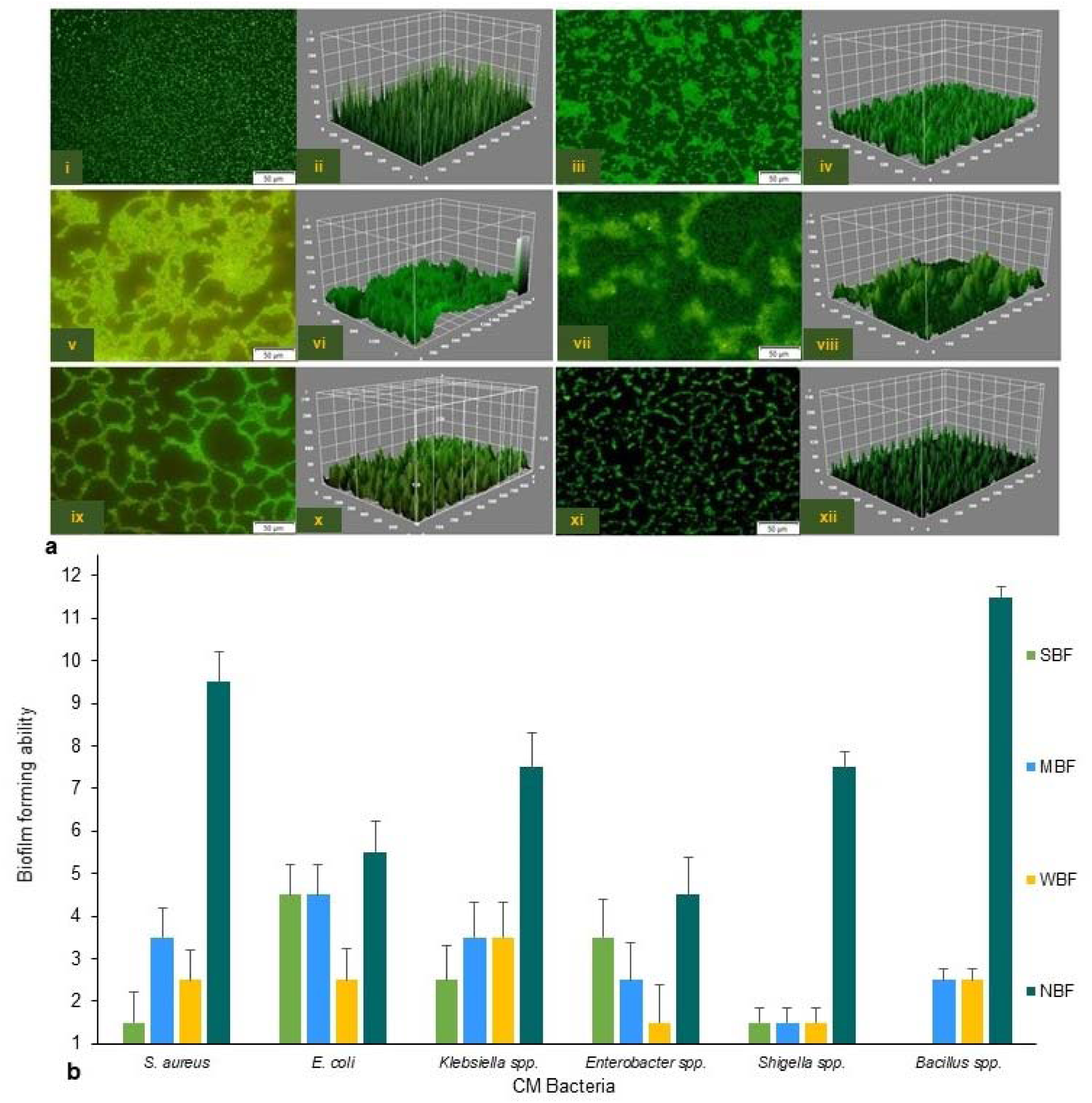
Biofilm formation (BF) ability of the six CM causing pathogens. BF assays was performed with solubilized crystal violet (CV) in a plate reader at 600 nm using 30% acetic acid in water as the blank and TSB as negative control. **a)** Confocal fluorescence images (2D and 3D) of *S. aureus* (i,ii), *E. coli* (iii,iv), *Klebsiella* spp. (v,vi), *Enterobacter* spp. (vii,viii), *Bacillus* spp. (ix,x) and *Shigella* spp. (xi,xii). Scale bars are indicated in μm. **b)** Category of the biofilm formation by six CM causing bacteria. The BF ability of the tested bacteria were classified as follows: NBF, non-biofilm formers optical density (OD) ≤ optical density cut-off (ODc); WBF, weak biofilm formers (ODc < OD ≤ 2 x ODc); MBF, moderate biofilm formers (2 x ODc < OD ≤ 4 x ODc), SBF, strong biofilm formers (OD > 4 x ODc). The ODc value was set as 0.045 and the mean OD of the negative control was 0.039±0.002. Thus, bacterial biofilms were divided into breakpoint categories; OD < 0.045 non-biofilm producers; OD ≥ 0.046 but ≤ 0.090 weak biofilm producers; ≥OD 0.091–≤0.180 moderate or partial biofilm producers; >0.181 strong biofilm producers. The results are presented as the mean ± SD, and post hoc Bonferroni test was used to compare the biofilm OD600 mean values (*p*<0.05).

### Pathogenic functional potentials genome of the CM microbiomes

We also investigated the possible links between chemotaxis and pathogenicity through the identification of putative genes or proteins associated with both flagellar motility and bacterial chemotaxis. The KEGG pathway analysis of MR tool identified 48 protein families associated with flagellar motility in prokaryotes, and among them, flagellar hook-length control protein, FliK (27.1%); flagellar biosynthesis proteins, FlhA, FliL, FliP, FlhF, FlgN, FliS, FlhB, FliO, FliQ (∼16.0%); flagellar M-ring protein, FliF (5.6%); and flagellar regulatory protein, FleQ (5.3%) were predominantly associated with cell motility (Supplementary Data 2). Twenty six functional genes encoding different proteins were found to be associated with bacterial chemotaxis (Supplementary Fig. 4, Supplementary Data 2), of them, methyl-accepting chemotaxis protein, mcp (44.2%); chemotaxis family proteins of bacterial two component system, CheV, CheA, CheB, CheBR, CheY (∼15.0%); aerotaxis receptor, Aer (7.5%); MotB (5.2%) and MotA (3.1%) were most abundant among these CM microbiotas (Supplementary Data 2). To explore the role of regulation and cell signaling mechanisms in mammary gland pathogenesis, using the SEED subsystem module of MR analysis, we found two-component regulatory systems BarA-UvrYBarA-UvrY(*sir*A) as the most abundant virulence regulatory gene (84.1%) in CM microbiomes (Supplementary Data 2). Another regulatory and cell signaling gene, endoplasmic reticulum chaperon *grp*78 (BiP) was also found as the single most abundant (93.8%) gene in proteolytic pathways of the CM associated bacterial strains (Supplementary Fig. 5, Supplementary Data 2). A deeper look at microbial genes associated with phages-prophages, transposable elements and plasmids revealed that pathogenicity islands related proteins such as methionine-ABC transporter substrate-binding protein (33.8%), GMP synthase (27.7%), tmRNA-binding protein; SmpB (16.0%), heat shock protein 60; GroEL (16.0%) and SSU ribosomal protein; S18p (6.1%) were predominantly abundant among the CM pathogens (Supplementary Data 2). The SEED module analysis also enabled us to identify 28 different protein functions associated with oxidative stress responses among the CM microbiomes which were mostly represented by catalase related proteins (26.7%), Cu-Zn-Fe-Mn mediated superoxide dismutases (12.7%), H_2_O_2_-inducible genes activator (7.8%) and paraquat-inducible protein B (7.3%) (Fig. 9, Supplementary Data 2).

**Fig. 9.**
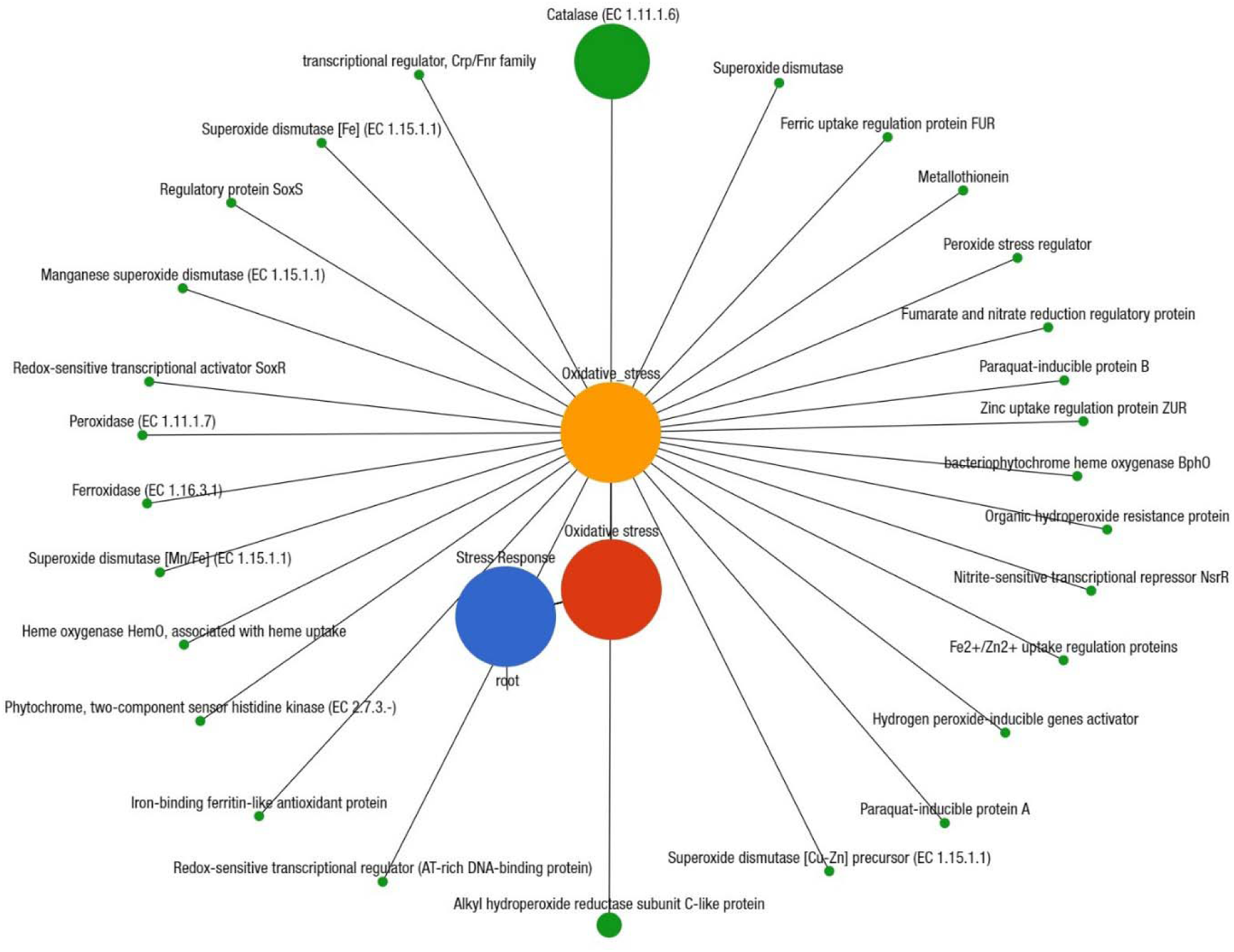
Projection of the clinical mastitis (CM) milk metagenome onto KEGG pathways. The whole metagenome sequencing (WMS) reveals significant differences (Kruskal–Wallis test, *p*=0.001) in functional microbial pathways. A total of 28 genes associated with oxidative stress were found in CM microbiomes. Black lines with green circles delineate the distribution of the stress related genes according to their class across the CM metagenome. The diameter of the circles indicates the relative abundance of the respective genes.

## Discussion

Previously, we reported that bovine CM milk microbiomes is a reservoir of diverse groups of resistiome (antibiotics and metal resistance, biofilm formation and quorum sensing genes) with functional biases in metabolism, bacterial chemotaxis, virulence regulation, compared to healthy milk microbiomes^2^. In this study, we employed a combination of both *in silico* (whole metagenome sequencing, WMS) and *in vitro* (culture-based) approaches to elucidate the resistome diversity in CM associated microbiomes. Recently, the WMS and other high-throughput sequencing (targeted amplicon) studies have provided new insights into the structure, function and dynamics of bovine CM milk^2–4,12^ and human lactational mastitis milk^24^ microbiomes. Our present findings are sufficiently enriched in taxonomic resolution and predicted protein functions, and corroborates to the findings of several previous studies^2–4,24^. The occurrence of bovine mastitis could be affected by cattle breeds^12,15,16^, and the diversity of CM-causing pathogens is associated with broad range of host-defense mechanisms as part of its immunological arsenal^25,26^. We found significant differences in taxonomic diversity and abundances among the CM microbiomes of four dairy breeds. The XHF cows suffering from CM had higher microbial diversity at strain-level, and a significant proportion of the microbiota found to be shared with that of the other three breeds (LZ, SW and RCC). Consistent with the results of earlier studies^12,15,16,26^, the taxonomic profile of the CM microbiomes found in four breeds of cows were dominated by phyla *Proteobacteria*, *Bacteroidetes*, *Firmicutes*, *Actinobacteria* and *Fusobacteria*. This breed specific variation in taxonomic richness and diversity of microbiome, especially in XHF and LZ cows, could be associated with their increased disease resistance or immune response^12,15,16^ and rumen microbial features (e.g., taxa, diversity indices, functional categories, and genes)^26^. However, further investigations will be necessary to evaluate the real effect of breed specific bacteria on cow mammary gland diseases.

Based on previously available culture-based reports on dairy animal mastitis pathogens in Bangladesh^6,27^ and other countries^1,7–9^, we identified six aerobic bacteria (*S. aureus*, *E. coli*, *Klebsiella*, *Enterobacter*, *Bacillus* and *Shigella*) through 16S ribosomal RNA (16S rRNA) gene sequencing and phenotypic characterizations, and these findings are in line with the taxonomic signature of WMS. Recent understanding regarding evolutionary relationships of major CM causing bacteria are primarily based on 16S rRNA gene phylogenetic identification along with a few individual gene or protein sequences^28^, which often produces conflicting phylogenies. This study also explored that the prevalence of CM milk pathogens could vary according to geographical locations and farming (semi-intensive to intensive grazing system in SER, semi-intensive to free-range grazing systems in CR) systems^1^. These differences may imply that the etiology of bovine CM in Bangladesh could be related to the breed/host genetic factors^12,15,16,26^, types of feeding and farm locations and types^1^, and types of antibiotics and/or metals used for treatment or other factors as have been described in other countries^1,8,9^.

Data presented here coupled with the data reported in our earlier study^2^ provides important insights into the diversity of resistomes in CM microbiomes. Our results are concordant with MDR bacteria reported elsewhere from the milk of clinically infected cows^8,15,21^, buffalo cows^9^ and humans^11,29^. Our findings linked multidrug resistance to efflux pumps (MREP), *Cme*ABC operon, *mdt*ABCD cluster, *Bla*R1 family, methicillin resistance in *Staphylococcus* (MRS), resistance to fluoroquinolones (RFL), and multiple metals resistance to CZCR and AR as the predominantly abundant antibiotics and toxic compounds resistance (RATC) functional groups in CM microbiomes suggesting that bovine CM milk microbiome constitutes a good reservoir for antimicrobial resistance^2,11, 29–33^. It has been reported that efflux pumps regulated by two-component systems in several pathogens, including *A. baumannii* and *K. pneumonia*, provide multidrug resistance, which may limit the treatment options against bacterial infections of the mammary glands^31,32^. Relative over-expression of efflux pumps enhances the resistances to antimicrobials by reducing the accumulation of antibiotics inside of the bacterial cells and providing sufficient time for the bacteria to adapt to the antibiotics (slow phase antibiotic efflux), and through mutations or alteration of antibiotic targets^31,33^. The *Cme*ABC operon is highly potent against multiple antibiotics, promotes the emergence of ARGs, and confers exceedingly high-level resistance to fluoroquinolones^33^. Therefore, multidrug resistance to efflux pumps and multiple heavy metals resistance represented ubiquitous resistance mechanisms among CM microbiomes, which might be associated with unethical overuse of antibiotics in dairy animals^8,9,15, 19–21^ and extensive application of toxic chemicals and metals in agricultural use^1,22,34^ or might have a function in the gut microbiome that is still unknown^13,29,35,36^. The RATC genes detected in this study are of particular interest because there is concern that the use of this class of antibiotics or metals in veterinary medicine, particularly for food animals, may contribute to the development of resistance to this class of antimicrobial options in human^29,35^.

*In-vitro* antibiogram of this study report higher prevalence of resistance to tetracyclines (tetracycline and doxycycline), quinolones (nalidixic acid), penicillins (ampicillin) and phenols (chloramphenicol), similar findings were observed in previous studies on bovine mastitis^8,9^. The AMR profile of bovine CM pathogens for different antimicrobials could vary according to the type and origin of bacteria^8–10^ and host-population such as bovine^8,21^ and bubaline cows^9^. Consistent with bacterial needs, heavy metals can be transformed (e.g., oxidized, reduced, methylated, or complexed) and used as a source of energy, terminal electron acceptors, or enzyme structural elements^34^. The highest abundance of CZCR genes among CM pathogens is mainly due to the presence of Co, Zn, and Cd detoxification systems^34^. Although the knowledge on uncontrolled spread of ARGs in bovine mastitis pathogens^8^ are increasing, but information on toxic compounds or heavy metal resistance is yet unavailable. In this study, heavy metals (Cr, Co, Ni and Cu) tested for antibacterial sensitivity showed good efficacy, although knowledge on their mode of action is limited. Thus, with the increase of MDR bacteria in CM, it is imperative that new biocidal and antimicrobial formulations are needed. The MIC and MBC tested metals revealed e□ective antimicrobial efficacies against a wide range of AMR pathogens^1,22,36^. We found that Cr and Co compounds had the highest antimicrobial efficacy (MIC) against all of the tested bacteria supported by several previous studies^22,37^. Furthermore, our present findings also suggested that the host genetic component in cattle breeds can significantly regulate the composition of the milk microbiome^12,15,16^, albeit not associated with resistomes profiles. Biofilm formation is an important virulence factor that may result in recurrent or persistent udder infections^38^ and treatment failure through increased resistance to antibiotics and protection against host defences^39^. The relative overexpression of genes encoding *lsr*ACDBFGE operon, biofilm adhesion biosynthesis (BAB), protein *Yjg*K cluster and quorum sensing: autoinducer-2 synthesis (QSAU2) in CM microbiomes is in accordance with several earlier reports^2,39,40^. In this study, the relative abundance of the predicted proteins for biofilms and quorum sensing (BF-QS) varied significantly among the selected six bacterial taxa. The BF and QS can be the strain specific or genetically linked traits, representing a selective advantage in pathogenesis of bovine CM^40^. BF can enhance proliferation of reactive oxygen and nitrogen species^34^ that can survive antibiotic treatment leading to the transfer of ARGs^41^. In this study, overall, 76.2% of the isolates were detected as biofilm formers, and their ability to producing biofilm varied significantly^38,39^. A large number of food spoilage and/or pathogenic bacteria, including *Enterococcus faecalis*, *Enterobacter* spp., *Pseudomonas* spp., *Klebsiella* spp., *S. aureus*, *E. coli*, *B. cereus*, and others, have already been associated with biofilms from dairy niches^22,23, 38–40^, which supports our current findings.

Bacterial chemotaxis mediated by flagellar activities^41^, and the flagella mediated virulence factors are found in many pathogenic species of bovine CM microbiomes, making them a potential target for new antibacterial therapeutics^41^. The intra- and interspecies cell-to-cell communication in bovine CM microbiomes were associated with 26 different genes, which might have vital roles in the early phase of mastitis for attachment to or entry into the udder tissues and virulence regulation^42^ and bacterial colonization in mammary tissues like other suitable sites^43^. The cheA-cheY two-component system mediated bacterial chemotaxis also facilitates the initial contact of bacteria with mammary gland epithelial cells and contribute to effective invasion^44^. The two-component signal transduction system BarA-UvrY regulates metabolism, motility, biofilm formation, stress resistance, virulence and quorum sensing in CM pathogens by activating the transcription of genes for regulatory small RNAs^45^. The up-regulation of genes coding for proteolytic activity, *grp78* (BiP) during host-pathogen interactions in CM is associated with endoplasmic reticulum (ER) stress which further triggers proteolytic activities to initiate the mechanism of pathogenesis and cell death^46^. Catalase activity is a marker of bovine mastitis, which plays a central role in milk redox control and increases markedly during the pathophysiology of bovine CM^47^. Our present findings corroborated with previous reports^47,48^ that an elevated oxidative stress mediated by catalase activity might have originated either from the mammary gland and/or bacterial cells. During the pathogenesis of bovine mammary gland, bacteria are not rapidly killed by the phagocytic activity of bovine macrophages; rather, they survive within macrophages during prolonged infection due to secretion of catalase and superoxide dismutases, which by degrading H_2_O_2_ inhibit ROS mediated killing mechanism of the host^47,48^.

## Conclusions

The bovine CM milk microbiomes harbor diverse groups of resistomes and other virulence factors. The diversity of resistomes positively correlated with the diversity of the microbial communities. The efflux pumps mediated multidrug resistance, methicillin, fluoroquinolones and beta-lactamase resistance, and multiple heavy metals (e.g., cobalt, zinc, cadmium, arsenic and chromium) resistance were the predominating in CM pathogens. Cattle breed is also a predominant factor for CM associated microbiome diversity, although resistome diversity does not affected by the breed specific microbiome signature. In bovine CM, biofilms may involve in colonizing the pathogens to udder tissues and teat canals, have an important role in antimicrobials resistance, resistant marker transfer and other virulence expression. Furthermore, flagellar movement and chemotaxis, regulation and cell signaling, phages-prophages, transposable elements, plasmids and oxidative stress had association with the pathophysiology of bovine CM. Therefore, accurate and timely identification of CM microbiome and its associated resistomes along with selection of proper therapeutic regimens will help improve the antimicrobials stewardship for prevention and control of bovine CM in Bangladesh.

## Methods

### Screening for clinical mastitis (CM) and sampling

We screened 260 quarter milk samples collected from 260 clinical mastitis (CM) affected cows belonging to 50 smallholding dairy farms in two geographical regions of Bangladesh (central region, CR=160; southeastern region, SER= 100) (Supplementary Fig. 1). The cows represented four different breeds, including local zebu (LZ), red Chattogram cattle (RCC), Sahiwal (SW), and crossbred Holstein Friesian (XHF) at their early stage of lactation (within 10-40 days post-calving). A screening test for CM was conducted using the California Mastitis Test (CMT^®^, Original Schalm reagent, ThechniVet, USA)^49^. Approximately 15-20 ml of milk from each cow was collected under aseptic conditions in a sterile falcon tube during the morning milking (8.00-10.00 am), and kept on ice (at 4°C) for transport to the laboratory for subsequent processing.

### Metagenomic DNA extraction and sequencing

Genomic DNA (gDNA) from 25 randomly selected CM samples was extracted by an automated Maxwell 16 DNA extraction platform using blood DNA purification kits (Promega, UK) following previously described protocols^2^. DNA quantity and purity were determined with NanoDrop (ThermoFisher, USA) by measuring 260/280 absorbance ratios. Sequencing libraries were prepared with Nextera XT DNA Library Preparation Kit^50^ and paired-end (2×150 bp) sequencing was performed on a NextSeq 500 machine (Illumina Inc., USA) at the George Washington University Genomics Core facility. Our metagenomic DNA yielded 596.74 million reads with an average of 23.87 million (maximum=39.75 million, minimum=8.89 million) reads per sample (Supplementary Data 1).

### Sequence reads preprocessing

The resulting FASTQ files were concatenated and filtered through BBDuk^2^ (with options k=21, mink=6, ktrim=r, ftm=5, qtrim=rl, trimq=20, minlen=30, overwrite=true) to remove Illumina adapters, known Illumina artifacts, and phiX. Any sequence below these thresholds or reads containing more than one ‘N’ were discarded. On average, 21.13 million reads per sample (maximum=36.89 million, minimum=4.71 million) passed the quality control step (Supplementary Data 1).

### Microbiome diversity and community analysis

The shotgun whole metagenome sequencing (WMS) data were analyzed using both mapping-based and assembly-based hybrid methods of PathoScope 2.0 (PS)^51^ and MG-RAST (MR), respectively^52^. In PS analysis, a ‘target’ genome library was constructed containing all bacterial sequences from the NCBI Database using the PathoLib module^51^. The reads were then aligned against the target libraries using the very sensitive Bowtie2 algorithm^53^ and filtered to remove the reads aligned with the cattle genome (bosTau8) and human genome (hg38) as implemented in PathoMap (−very-sensitive-local -k 100 --score-min L,20,1.0). Finally, the PathoID^54^ module was applied to obtain accurate read counts for downstream analysis. In these samples, 17.20 million reads (4.3% of total reads) mapped to the target reference genome libraries after filtering the cow and human genome (Supplementary Data 1). The raw sequences were simultaneously uploaded to the MR server (release 4.0) with proper embedded metadata and were subjected to the quality filter containing dereplication and removal of host DNA by screening for taxonomic and functional assignment. Alpha diversity (diversity within samples) was estimated using the observed species, Chao1, ACE, Shannon, Simpson and Fisher diversity indices^55^ for both PS and MR read assignments and counts. To visualize differences in bacterial diversity, a principal coordinate analysis (PCoA) was performed based on weighted-UniFrac distances (for PS data) through Phyloseq R package (version 3.5.1)^56^ and Bray-Curtis dissimilarity matrix^57^ (for MR data). We have also used OmicCircos (version 3.9)^58^ which is an R package based on python script for circular visualization of both microbiome diversity and resistance to antibiotics and toxic compounds (RATC) functional groups found in MR data for respective four breeds of CM cows.

### *In vitro* identification of bacteria

Collected CM milk samples (n=260) were subjected to selective isolation and identification of *S. aureus*, *E. coli*, *Klebsiella*, *Enterobacter*, *Shigella* and *Bacillus* species according to previously described microbiological methods^1,6–9^. The pathogens were identified based on their colony morphology, hemolytic patterns on blood agar and Gram-staining^8^. Gram-positive bacteria were further confirmed based on their biochemical characteristics in indole, methyl red, Voges-Proskauer (VP), catalase, oxidase, urease and triple sugar iron (TSI) tests, and growth on mannitol salt agar. Gram-negative bacteria were confirmed based on the results of indole, methyl red, citrate (IMViC) tests and lactose fermentation on Mac agar^9,40^. Finally, all isolates were stored at −80 °C for further genomic identification.

### PCR amplification and ribosomal (16S rRNA) gene sequencing

Genomic DNA of probable *S. aureus*, *E. coli*, *Klebsiella*, *Enterobacter*, *Shigella*, and *Bacillus* species was extracted from overnight cultures using the boiled method^59^. The quantity and purity of the extracted DNA was determined as mentioned before. The 16S rRNA gene was amplified using universal primers 27F (5′-AGAGTTTGATCCTGGCTCAG-3′) and U1492R (5′-CTACGGCTACCTTGTTACGA-3′). Agarose gel electrophoresis (1.2% wt/vol) was used to verify the presence of PCR products. DNA sequencing was carried out at First Base Laboratories Sdn Bhd (Malaysia) using Applied Biosystems highest capacity-based genetic analyzer (ABI PRISM^®^ 377 DNA Sequencer) platforms with the BigDye^®^ Terminator v3.1 cycle sequencing kit chemistry^61^.

### Phylogenetic analysis of the microbial communities

Taxonomic abundance of the WMS data was determined by applying the ‘‘Best Hit Classification’’ option in PS pipeline using the NCBI database as a reference with the following settings: maximum e-value of 1×10^−30^; minimum identity of 95% for bacteria, and a minimum alignment length of 20 as the set parameters. A phylogenetic tree consisting of the top 200 abundant bacterial strains identified through PS analysis from the WMS reads of the 25 CM samples with >90% taxonomic identity was constructed using maximum-likelihood method in Clustal W (version 2.1)^61^ and visualized using interactive Tree Of Life (iTOL)^62^. Another, phylogenetic tree consisting of 40 strains correspondent to *in vitro* examined six CM bacteria found in 260 CM samples with >90% taxonomic identity was also constructed using same methods. Using Molecular Evolutionary Genetics Analysis (MEGA) version 7.0 for bigger datasets^63^, the 16S rRNA gene sequences, amplified from all individual bacterial isolates, were aligned with each other and with relevant reference sequences obtained from the NCBI Database, and a maximum-likelihood tree was generated using these 16S rRNA gene sequences^63^. The percentage of replicate trees in which the associated taxa clustered together in the bootstrap test (1000 replicates) is shown next to the branches^64^.

### Antimicrobial susceptibility testing

The *in vitro* antibiogram profile of 221 CM isolates was determined using the disk diffusion method following the Clinical Laboratory Standards Institute^65^ guidelines. Antibiotics were selected for susceptibility testing corresponding to a panel of antimicrobial agents (Oxoid™, Thermo Scientific, UK) of interest to the dairy industry and public health in Bangladesh. The selected groups of antibiotics were commonly used in treating CM by the dairy farmers and included penicillins (ampicillin, 10 μg/mL), tetracyclines (doxycycline, 30 µg/mL; tetracycline, 30 µg/ML), nitrofurans (nitrofurantoin, 300 µg/mL), quinolones (ciprofloxacin, 10 µg/mL; nalidixic acid, 30 µg/mL), cephalosporins (cefoxitin, 30 µg/mL), penems (imipenem, 10 µg/mL), phenols (chloramphenicol, 30 µg/mL), aminoglycosides (gentamycin, 10 µg/mL; vancomycin, 30 µg/mL), macrolides (erythromycin, 15 μg/mL). Resistance was defined according to CLSI (2017) with slight modifications^8,9^.

### Metal susceptibility testing

The antibacterial effect of heavy metals was evaluated *in vitro* for the isolated pathogens using both agar well diffusion and tube dilution methods^1,22^. Five heavy metals such as copper (Cu), zinc (Zn), chromium (Cr), nickel (Ni), and cobalt (Co) were used as salts: CuSO4.5H2O, ZnSO4.7H2O, K2Cr2O7, NiCl2, and CoCl2.6H2O, respectively to study the level of zone of inhibition (ZOI). Briefly, pure culture of the isolated pathogens from NA plates were sub-cultured into Mueller-Hinton agar (Oxoid^TM^, UK) plates, and five 7 mm wells were made, one in the center of the plate and the other four about 20 mm away from the center. Varying concentrations of the metal solutions were prepared (2, 4, 8, 16, 32, 48 and 64 μ mL) and 100μ of prepared solution was inoculated into the central well of 1 cm in diameter. The plates were incubated at 37 °C for 24 h to allow diffusion of the metal into the agar, and the antibacterial activity was determined by measuring the diameter of ZOI in mm^12^. After investigating the resistance profile of the isolates at different concentrations, the minimal inhibitory concentration (MIC) of the metals was determined by the tube dilution method by gradually increasing or decreasing the heavy metal concentrations^1^. Finally, growth of bacterial colonies was observed and the concentration that showed no growth was considered as the minimum bactericidal concentration (MBC)^1^.

### Biofilm assay and microscopy

Microtiter plate assays were performed to screen for biofilm formation (BF) ability of 80 randomly selected isolates using standard protocols^22,23,38,39^. We quantified the absorbance of solubilized crystal violet (CV), in a plate reader at 600 nm using 30% acetic acid in water as the blank and TSB as negative control. The solution was removed, and the absorbance measured at optical density-590 (OD590) (n = 3). To determine BF ability of strains, cut-off OD (ODc) was defined as three standard deviations above the mean OD of the negative control. Strains were classified as: non-biofilm formers, NBF (OD ≤ ODc); weak biofilm formers, WBF (ODc < OD ≤ 2 x ODc); moderate biofilm formers, MBF (2 x ODc < OD ≤ 4 x ODc) and strong biofilm formers, SBF (OD > 4 x ODc)^22,39^. In this study, the ODc value was set as 0.045 and the mean OD of the negative control was 0.039±0.002^22^. The biofilms were then visualized using 5% TSB as nutrient rich media and FilmTracer™ LIVE/DEAD^®^ Biofilm Viability Kit as staining materials under Olympus BX51 upright microscope at 40X objective, and finally images were collected using Olympus DP73 camera through cellSens entry software (Olympus Corporation, Japan) and visualized using image J software^39^. As a negative control, we used *E. coli* DH5 alpha for all the *in vitro* resistome (antimicrobial and metal susceptibility tests and biofilm assays) analysis tests.

### Microbial functional analysis

Metagenomic functional composition was based on the gene families from different levels of SEED module and the Kyoto Encyclopedia of Genes and Genomes (KEGG) database^66^ using the MG-RAST 4.1 (MR) pipeline^52^. We observed significant differences (Kruskal–Wallis test, *p*=0.001) in the relative abundance of genes coding for RATC and microbial functional genomic potentials in four cattle breeds.

### Statistical analysis

The characteristics of breeds of the cows with CM were compared using a Kruskal–Wallis test for quantitative variables^2^. The Shapiro-Wilk test was used to check normality of the data, and the non-parametric test Kruskal-Wallis rank sum test was used to evaluate differences in the relative abundance of bacterial taxa at strain level according to breed groups^12,15,16^. The statistical analyses for the MR data were initially performed by embedded calls to statistical tests in the pipeline and validated further using IBM SPSS (SPSS, Version 23.0, IBM Corp., NY USA) using the above mentioned tests. For the functional abundance profiling, the statistical (Kruskal–Wallis test and Pearson correlation) tests were applied at different KEGG and SEED subsystem levels in the MR pipeline^52^. To evaluate the significant relationships between identified bacterial species and the study region, we used the two-sample proportions test using SPSS. Results were considered statistically significant when *p*<0.05 and highly significant when *p*<0.01. Mean values were used to compare the antimicrobial efficacy results of the tested antibiotics and heavy metals at varying concentrations. Standard error means were calculated to analyze the distributions of the data from the mean value and confidence intervals of 95% were calculated for the MIC and MBC tests results to plot error bars^22,39^. We also performed Pearson correlation tests to test for relationships between taxonomic abundance of the pathogens and antimicrobial resistance both for cultural and metagenomic data. A post hoc Bonferroni test was used to compare the biofilm OD600 mean values^22,39^.

## Acknowledgements

The authors thank Stephanie Warnken, PhD student at the Computational Biology Institute, Milken Institute School of Public Health, The George Washington University, USA for their technical support in learning basic bioinformatics operations. We also acknowledge the cooperation of the dairy farmers for allowing us to conduct the study in their farms.

## Author contributions

M.N.H., M. S., A.I. and M.A.H. conceived and designed the overall study. M.N.H. surveyed and collected all the field samples. M.N.H., R.A.C., K.M.G, O.S. and O.K.I. carried out laboratory works including DNA extractions and sequencing, microbiological (cultural, biochemical) examinations, antimicrobial (antibiotics, metals) sensitivity tests and biofilm assays. M.A.H. and K.A.C. contributed chemicals and reagents. M.N.H. and A.I. conceived, designed and executed the bioinformatics analysis. M.N.H. interpreted the results and drafted the manuscript. M.S., K.A.C. and M.A.H contributed intellectually to the interpretation and presentation of the results. Finally, all authors have approved the manuscript for submission.

## Supplementary Information

Supplementary information supporting the findings of the study are available in this article as Supplementary Data files, or from the corresponding author on request.

## Data availability

The sequence data reported in this paper have been deposited in the NCBI database (BioProject PRJNA529353 for metagenome sequences, NCBI accession number: MN 620423-MN 620430 for 16S rRNA gene sequences) and are available from the corresponding author upon reasonable request.

