## Supplementary figures and images for "Resistome diversity in bovine clinical mastitis microbiome, a signature concurrence"

### Supplementary Fig. 1

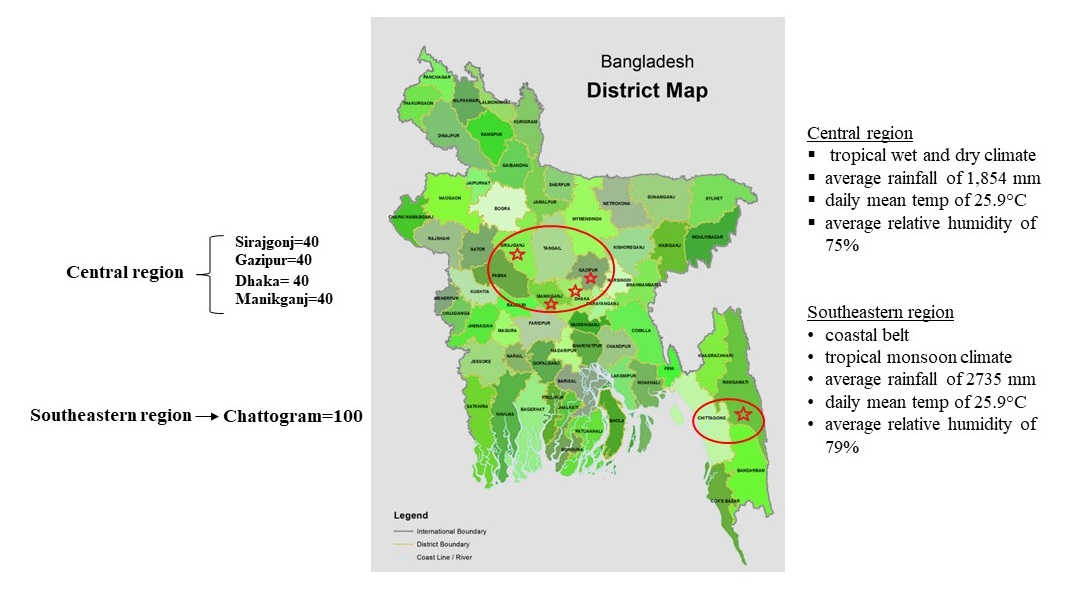

### Supplementary Fig. 2

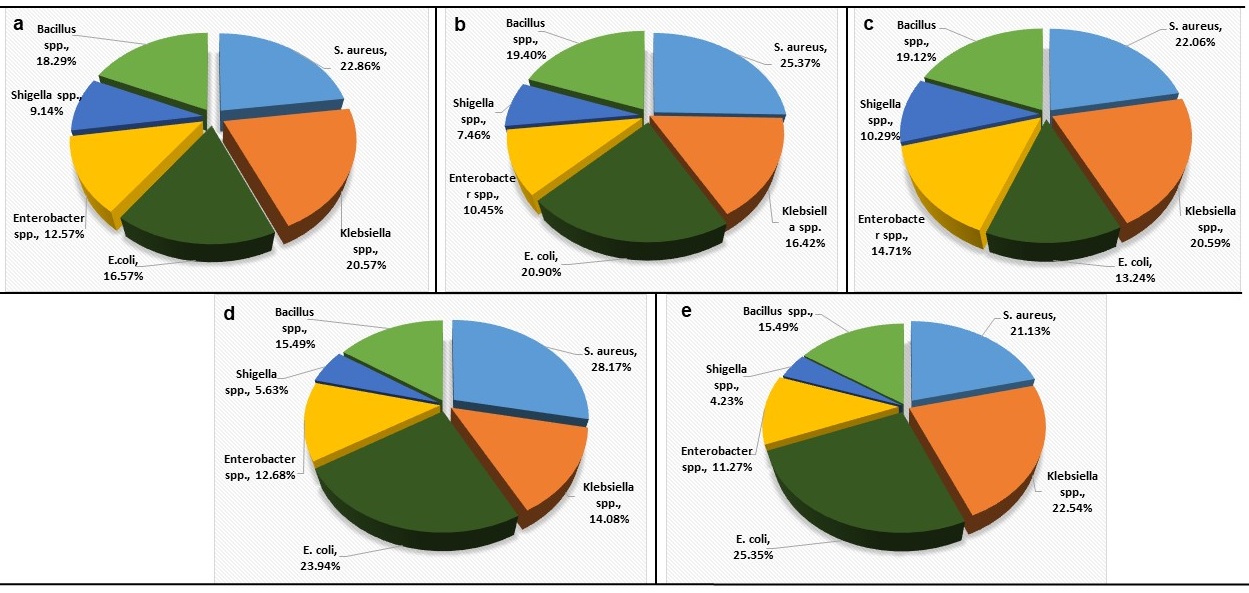

### Supplementary Fig. 3

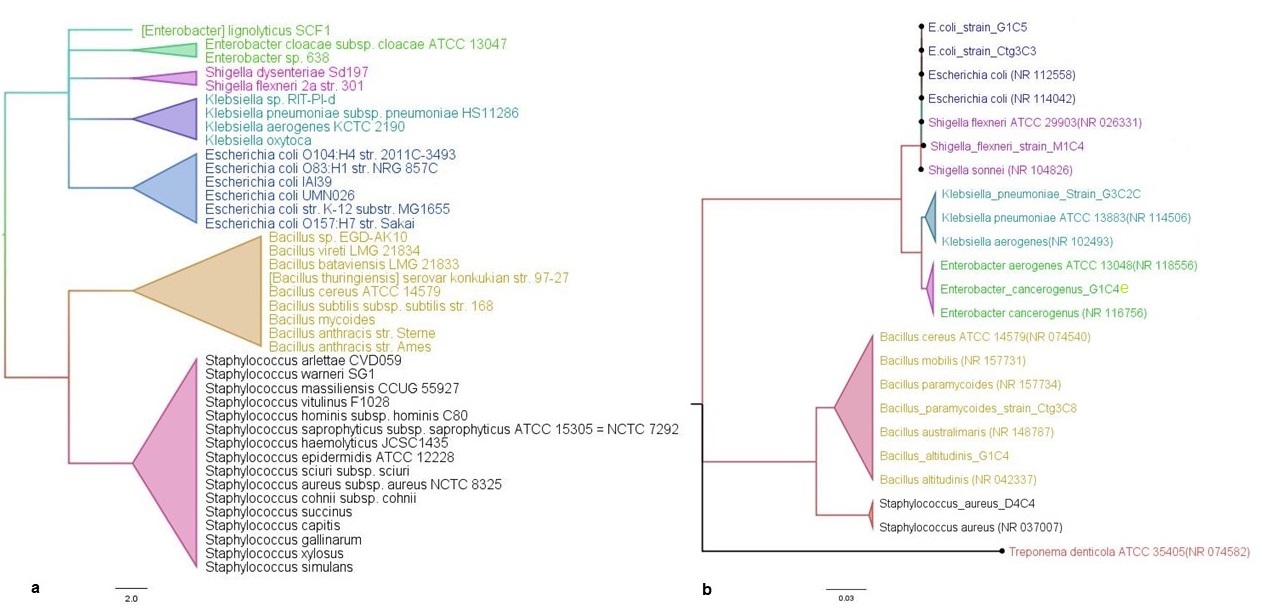

### Supplementary Fig. 4

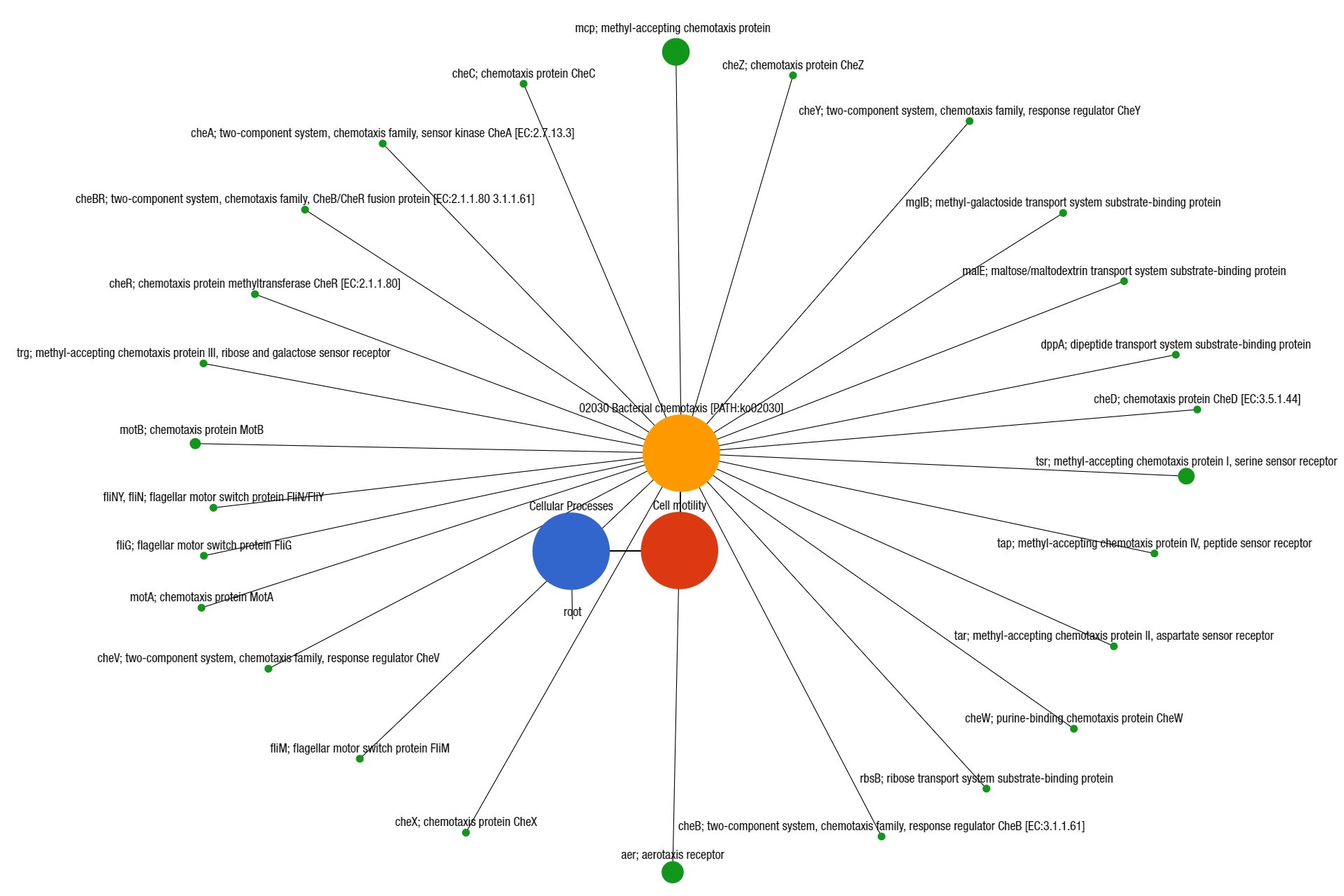

### Supplementary Fig. 5

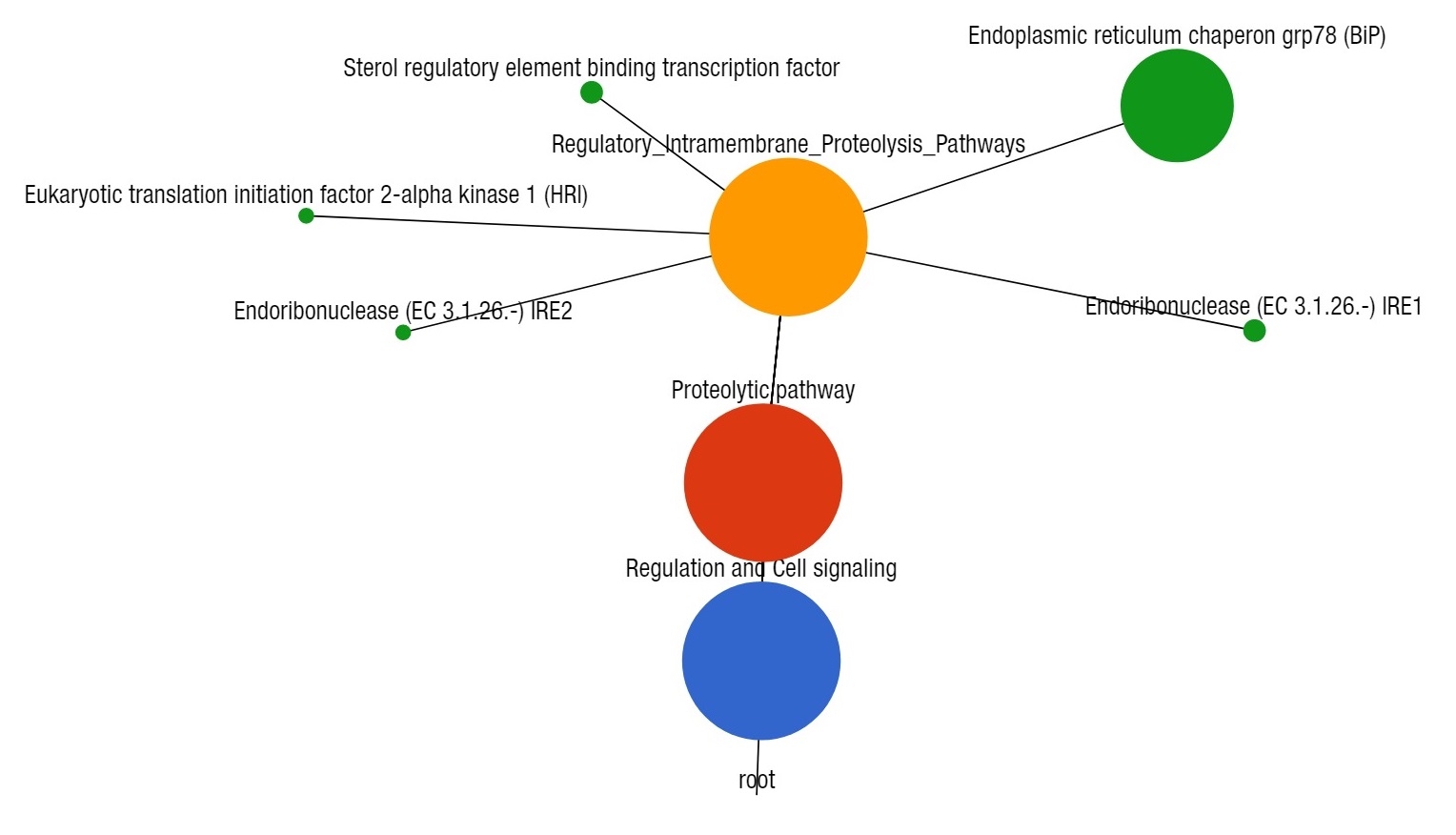
