## Supplementary Fig. legends for "Resistome diversity in bovine clinical mastitis microbiome, a signature concurrence"

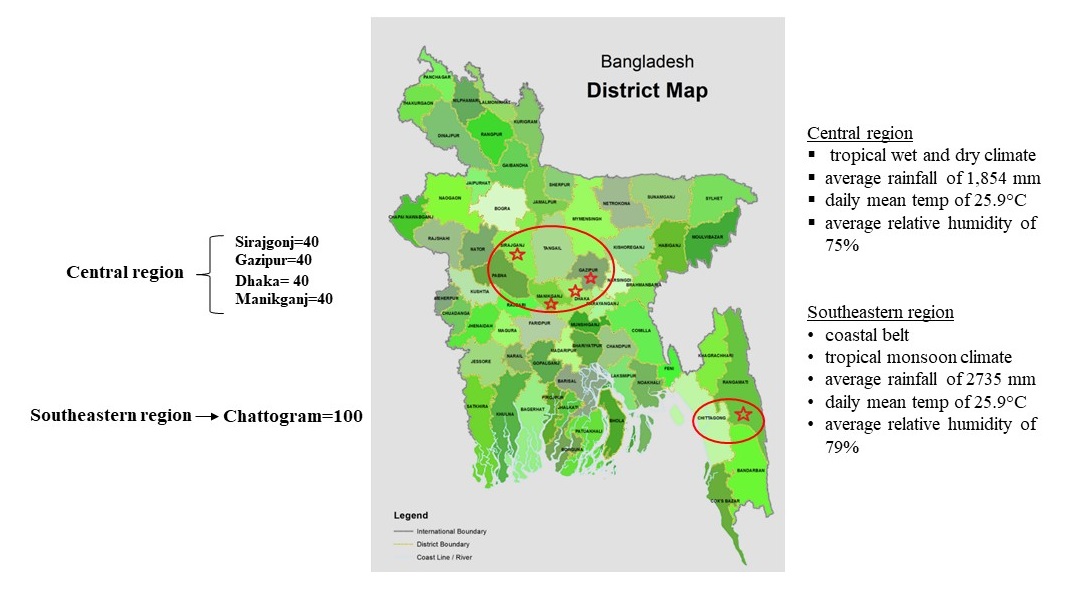

**Supplementary Fig. 1** Study areas. This study was conducted in two different regions of Bangladesh. The lower red circle indicates the southeast coastal region (SER); Chattogram district, and the top red circle represents the central region (CR) which includes four different (Dhaka, Gazipur, Manikgonj and Sirajgonj) districts of Bangladesh. A total of 260 CM samples (SER, 100; CR, 160) were collected from four different dairy breeds.

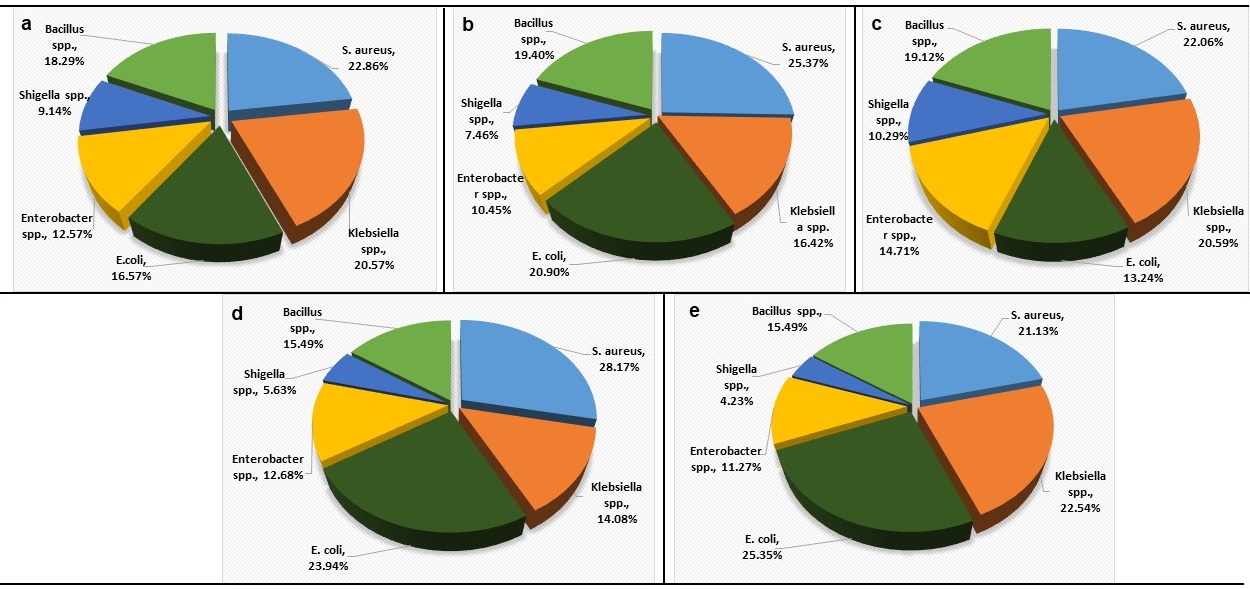

**Supplementary Fig. 2** Genera/species-level prevalence and distribution of six bacteria isolated from bovine CM milk. Microorganisms identified in bovine CM samples originated from two different regions **a**, southeastern region; **b-e**, central region) of Bangladesh. The southeastern region (SER) included 175 isolates and the central region (CR) included 277 isolates. The prevalence of the identified microorganisms varied significantly (*p*=0.01) between (SER, CR) and within (b-e) the study regions. This study demonstrated *S. aureus* as the chief etiology of bovine CM in Bangladesh while *Shigella* species remained as the least frequently detected bacterial pathogen causing mammary gland inflammation of the dairy cows.

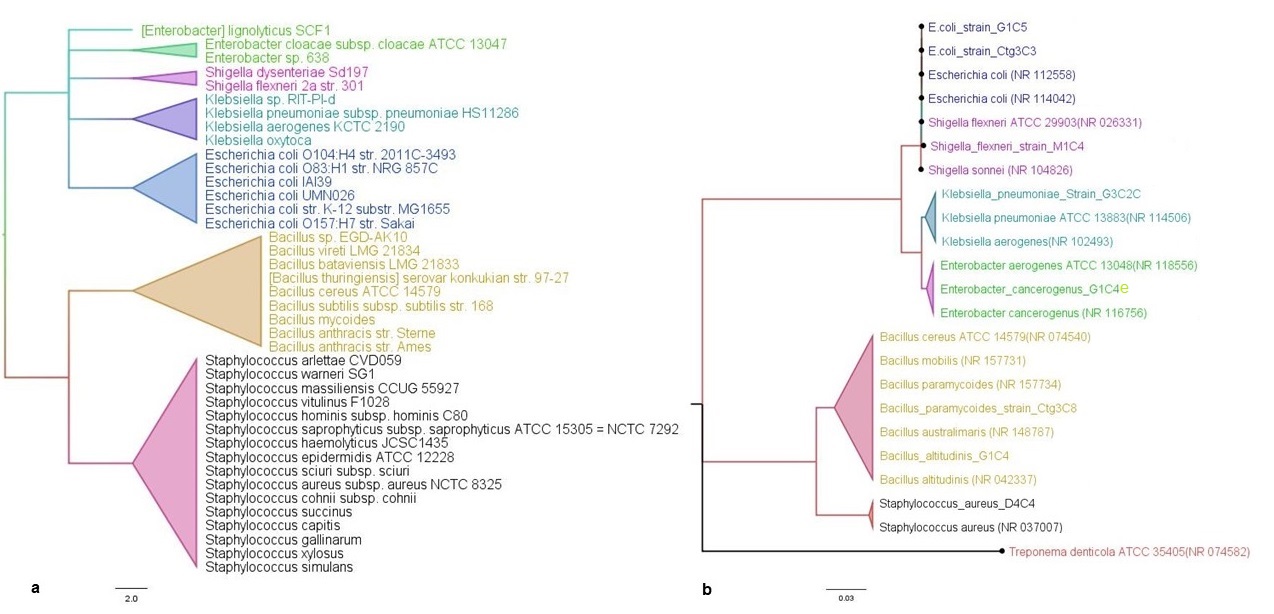

**Supplementary Fig. 3** Phylogenetic diversity of the bovine clinical mastitis (CM) pathogens. **a** consensus tree based on alignment of the whole metagenome sequencing (WMS) data. **b** consensus tree based on alignment of the ribosomal (16SrRNA) gene sequencing. The first phylogenetic tree was constructed with >80% taxonomic identity using the neighbor-joining method in Clustal W while the second one in MEGA, and both trees were visualized through FigTree. The same bacterial species and/strains identified in both WMS and 16S sequencing have presented with same color codes such as *S. aureus* in black, *Bacillus* species in deep yellow, *Enterobacter* species in green, *Klebsiella* species in paste, *Shigella* species in pink and *E. coli* in blue colors. The scale bars indicate 2.0 and 0.02 substitutions per nucleotide position respectively in WMS and 16S gene sequences. More details about the relative abundance of the identified strains in WMS can be found in the text and in Supplementary Data 1.

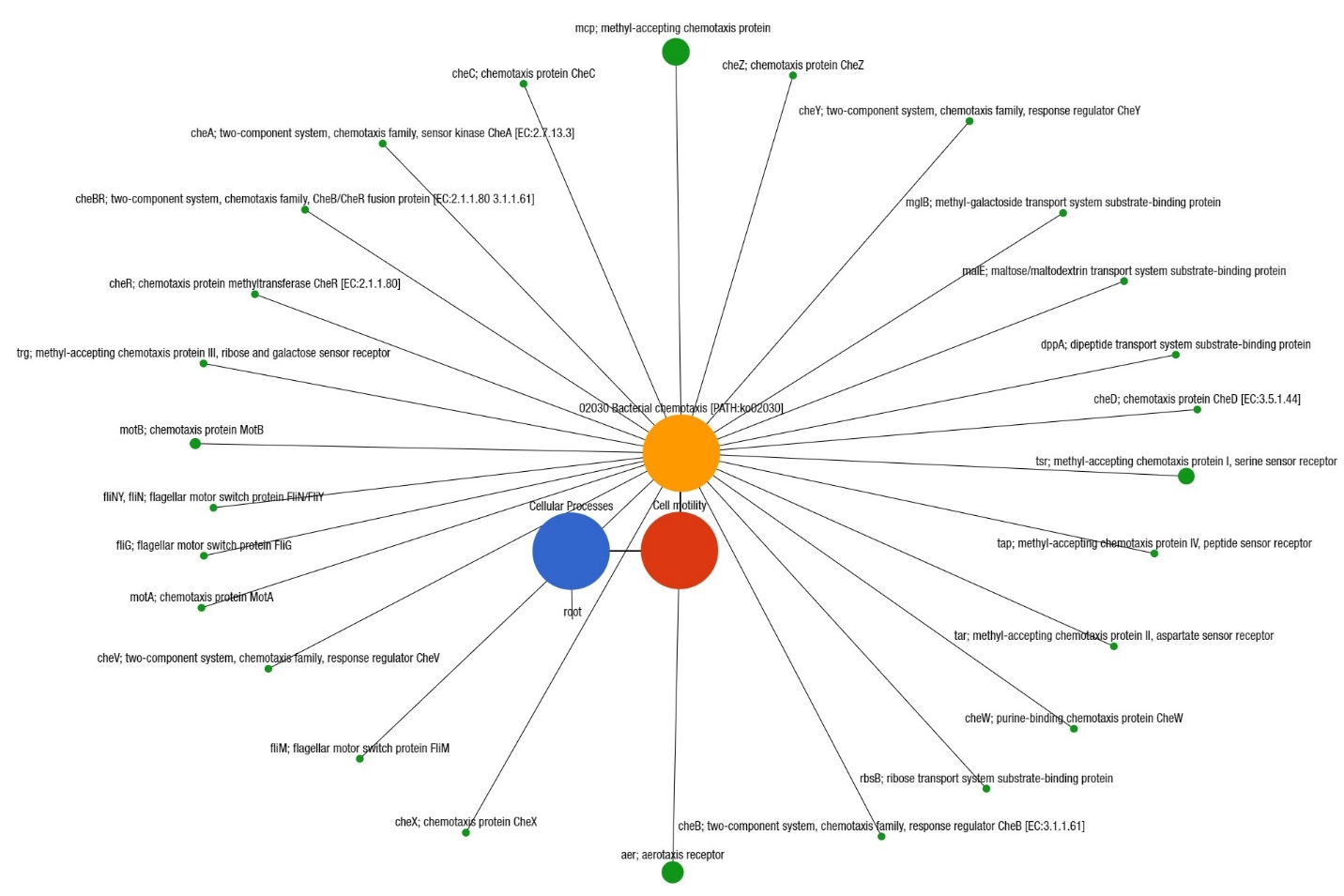

**Supplementary Fig. 4** Projection of the clinical mastitis (CM) milk metagenome onto KEGG pathways. The whole metagenome sequencing (WMS) reveals significant differences (Kruskal–Wallis test, *p*=0.001) in functional microbial pathways. A total of 26 genes associated with bacterial chemotaxis were found in CM microbiomes. Black lines with green circles demarcate the distribution of the chemotaxis related genes across the CM milk samples. The diameter of the circles indicates the relative abundance of the respective genes.

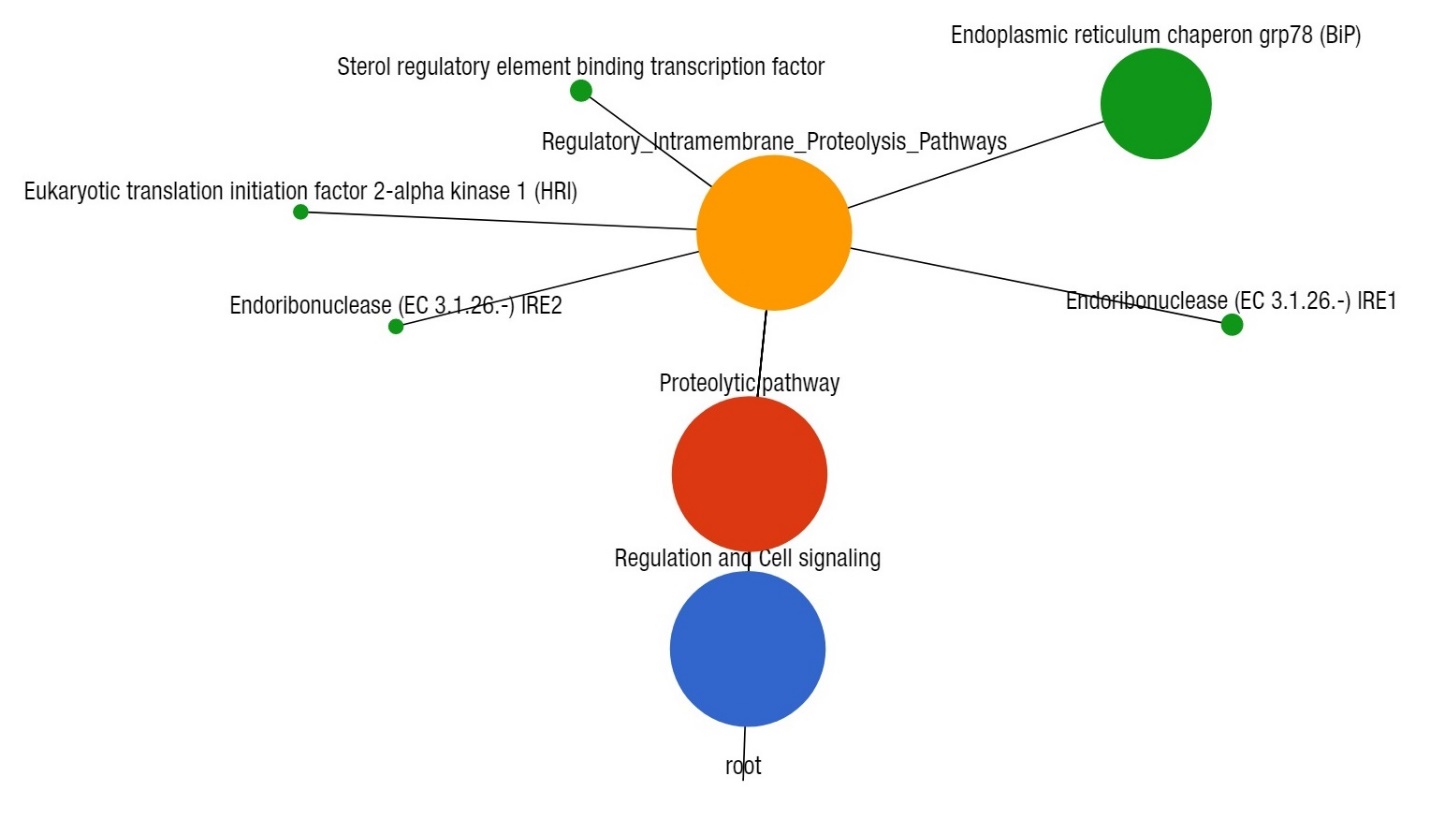

**Supplementary Fig. 5** Projection of the clinical mastitis (CM) milk metagenome onto KEGG pathways. The whole metagenome sequencing (WMS) reveals significant differences (Kruskal–Wallis test, *p*=0.001) in functional microbial pathways. Five genes associated with proteolytic activities were found in CM microbiomes. Black lines with yellow circles demarcate the distribution of the resistant genes according to their class across the metagenome. The diameter of the circles indicates the relative abundance of the respective genes.
