## Supplementary Table 1 for "Resistome diversity in bovine clinical mastitis microbiome, a signature concurrence"

| Farm location | *Staph. aureus* | | *E. coli* | | *Klebsiella* spp. | | *Enterobacter* spp. | | *Shigella* spp. | | *Bacillus* spp. | |
| --- | --- | --- | --- | --- | --- | --- | --- | --- | --- | --- | --- | --- |
|  | Sample | Isol.^1^ | Sample | Isol.^1^ | Sample | Isol.^1^ | Sample | Isol.^1^ | Sample | Isol.^1^ | Sample | Isol.^1^ |
| Chattogram | 24 | 40 | 16 | 29 | 20 | 36 | 12 | 22 | 10 | 16 | 18 | 32 |
| Dhaka | 10 | 17 | 8 | 14 | 7 | 11 | 4 | 7 | 4 | 5 | 7 | 13 |
| Gazipur | 8 | 15 | 5 | 9 | 8 | 14 | 6 | 10 | 5 | 7 | 8 | 13 |
| Manikgonj | 11 | 20 | 9 | 17 | 6 | 10 | 5 | 9 | 3 | 4 | 6 | 11 |
| Sirajgonj | 8 | 15 | 10 | 18 | 9 | 16 | 5 | 8 | 2 | 3 | 6 | 11 |
| Total | 61 | 107 | 48 | 87 | 50 | 87 | 32 | 56 | 24 | 35 | 45 | 80 |

Supplementary Table 1. Distribution of 452 bacteria isolated from 260 clinical mastitis (CM) sample in 50 smallholding dairy farms of Bangladesh
