## Supplementary Table 2 for "Resistome diversity in bovine clinical mastitis microbiome, a signature concurrence"

| Tested metals | *S. aureus* | | *E. coli* | | *Klebsiella* sp. | | *Enterobacter* sp. | | *Shigella* sp. | | *Bacillus* sp. | |
| --- | --- | --- | --- | --- | --- | --- | --- | --- | --- | --- | --- | --- |
|  | MIC | MBC | MIC | MBC | MIC | MBC | MIC | MBC | MIC | MBC | MIC | MBC |
| Cu | 33.21±1.1 | 38.44±0 | 29.11±0 | 36.22±1.8 | 21.42±0 | 31.2±0 | 23.10±0 | 25.60±2.5 | 7.5±1.9 | 25.12±0 | 19.0±0.5 | 35.12±2.1 |
| Zn | 38.1±0.9 | 41.23±1.2 | 19.21±0 | 24.21±1.3 | 22.4±0.2 | 30.2±2.3 | 19.72±0 | 25.14±1.3 | 7.5±1.8 | 33.11±0 | 27.1±0 | 41.21±1.4 |
| Cr | 13.2±0 | 11.32±0.7 | 7.42±0 | 13.02±0 | 5.82±0.8 | 12.02±0 | 9.4±0 | 25.52±0.8 | 3.4±0.7 | 15.82±0 | 9.3±0 | 19.44±0 |
| Co | 8.74±0.6 | 21.32±0 | 10.4±0.6 | 14.3 ±0.4 | 7.8±1.2 | 19.33±2.3 | 17.2±0 | 19.2±0 | 5.0±0.5 | 27.02±0.8 | 15.3±0 | 15.28±0.2 |
| Ni | 20.1±0.9 | 23.1±2.2 | 28.21±0.2 | 35.11 ±1.2 | 26.8 ±1.6 | 34.0±1.3 | 22.12±0 | 24.45±0 | 3.5±1.4 | 28.12±0 | 19.3±0.5 | 31.33±0.7 |

Supplementary Table 2. Minimal inhibitory concentration (MIC) and minimal bactericidal concentration (MBC) values (μg/mL) for the metals against tested six bacteria demonstrating that chromium and cobalt displayed the most inhibitory concentrations (MIC), and cobalt, chromium and nickel demonstrated the most potent MBCs.

Cu=copper, Zn=zinc, Cr= chromium, Co=cobalt, and Ni=nickel.
